# Prior knowledge reveals two computational regimes for syntactic processing in the human brain

**DOI:** 10.64898/2026.07.11.737945

**Authors:** Cosimo Iaia, Alessandro Tavano

## Abstract

The human brain rapidly transforms continuous speech into structured, meaningful linguistic representations, yet how prior knowledge constrains this process remains unclear. To characterize this influence, we combined MEG recordings acquired during audiobook listening with corpus-derived transition probabilities over syntactic features defined within both phrase-structure and dependency-based grammars. Prior knowledge enhanced the neural representation of memory-related features indexing syntactic structures that must remain open for future completion, an effect that generalized across grammatical formalisms. This enhancement was strictly local: only immediately preceding contexts improved neural decoding at both word onset and offset, whereas longer histories either returned decoding performance to baseline at the word level or impaired decoding. By contrast, integration-related features indexing the completion of syntactic operations showed no benefit from prior knowledge and were represented most strongly at word offset, consistent with structural resolution being implemented as a word-level operation. These dissociable dynamics reveal two concurrent neural computational regimes for syntactic processing: a forward-looking, locally maintained predictive code for pending structure and a backward-looking, integrative code engaged when structure is resolved.

## 1 Introduction

The human brain continuously generates expectations about upcoming linguistic input, a process often formalised in terms of word surprisal [1–7]. Indeed, it has been proposed that humans and Large Language Models (LLMs) share next-word prediction as a fundamental cognitive process [8]. However, cognitive similarities between humans and machines are difficult to evaluate because, despite the well-established predictive nature of speech and language processing, existing neural decoding approaches primarily characterize word-level (current) syntactic state, rather than the *expectations* that listeners have about how sentence structure will develop [9]. Native speakers acquire strong priors over sentence structure through language acquisition mechanisms [10] and lifelong exposure to the statistical regularities of language use [1, 2]. Behavioral evidence strongly supports the role of prior knowledge in shaping perceptual and linguistic expectations [1, 2]. Such priors can also sharpen the neural correlates of perceptual representations [11, 12]; however, comparable neural evidence is still lacking for the internally generated structure-building operations of human language. To approximate the effects of a lifelong language prior, we used the Brown corpus [13] together with a Markov chain approach and estimated the probability of each word’s syntactic feature from up to five preceding items.

We specifically targeted syntactic operations essential for online comprehension, that is, the dynamic integration of incoming words into an evolving syntactic representation. Sentence perception unfolds along at least two concurrent dimensions: the linear sequence of words and the hierarchical syntactic structure constructed over them [14–17]. These dimensions place distinct demands on working memory, which Dependency Locality Theory formalises in terms of costs: integration costs arise when an incoming word is linked to previous material - sometimes a few words back-, whereas memory costs arise when incomplete syntactic relations are actively maintained until they can be resolved via structure building operations [18, 19].

The extensive empirical work on memory and integration costs mostly relies on dependency grammar notation [20], in which logico-semantic relationships between words are directly represented as labeled dependencies, and structure is inferred. Dependency-based grammars are important as they enable scalable, high-performing large-scale text analysis and information extraction [21–27]. However, neural investigations of continuous speech typically rely on syntactic measures derived from phrase structure notation, in which tree configurations are explicit, and semantic relationships are structurally mediated. To bridge the Natural Language Processing (NLP) and neuroscience communities around both phrase-structure and dependency grammars as powerful algorithmic hypotheses [28–30] we indexed potential changes in syntactic structure held in memory [9, 31, 32], and captured relation integrating operations [33–35] using both grammatical notations. To characterize the precise dynamics of prior knowledge effects, we resorted to generalized additive mixed models (GAMMs [36]), which enabled us to test nonlinearities in decoding profiles across contexts and feature types.

We found that memory and integration operations have dissociable neural decoding profiles, with stronger onset-related effects for memory measures and stronger offset-related effects for integration measures, thereby justifying their theoretical construct. Prior knowledge significantly improved the neural decoding of memory but not of integration measures. Crucially, this benefit followed an inverted-U profile, extending no further than the two to three preceding items before tapering off or even decreasing. Human language, in this view, is not an engine of unbounded contextual accumulation, but a nonlinear system that builds structure online under tight locality constraints [37].

## 2 Results

### 2.1 Decoding syntactic states

We first derived and validated phrase-structure and dependency grammar metrics to quantify the current syntactic weight assigned to each word (baseline decoding). Phrase-structure grammars represent syntactic relations as hierarchical trees of lexical items and constituents, yielding a node-based structure in which terminal nodes correspond to words and non-terminal nodes denote constituents labeled by their syntactic functions (e.g., noun phrase, verb phrase) (Fig. 1 A, middle panel). For each word, we extracted the number of closing nodes [38], the number of open nodes [9, 31], and the depth of tree structure [9]. We also derived a variant of tree depth in which a value of +1 is added to the depth level of each word when a phrase boundary is detected (which we labeled *last right*, see Methods). In dependency grammars, logico-syntactic relations are represented as labeled dependencies between words (Fig. 1 A, lower panel). For each word, we computed the (absolute) dependency distance (or dependency length) [39], the directed dependency distance by maintaining positive and negative signs to the syntactic weights, and the dependency depth. These measures were grouped into memory measures (*tree depth, last right, dependency depth*, and *open nodes*) and integration measures (*absolute dependency distance, dependency distance*, and *closing nodes*) (Fig. 1 B, middle panel and lower panel, respectively). Memory measures tag the syntactic representations that have to be maintained in memory as they provide one or more structural positional options for the upcoming linguistic units, while integration measures indicate the insertion of one or more syntactic items into the preceding structure to close an existing position. We then fit a liner decoding model for each feature independently for each feature and each time point on an openly available MEG data set of participants listening to stories (see Methods, see Fig. 1 D).

**Fig. 1.**
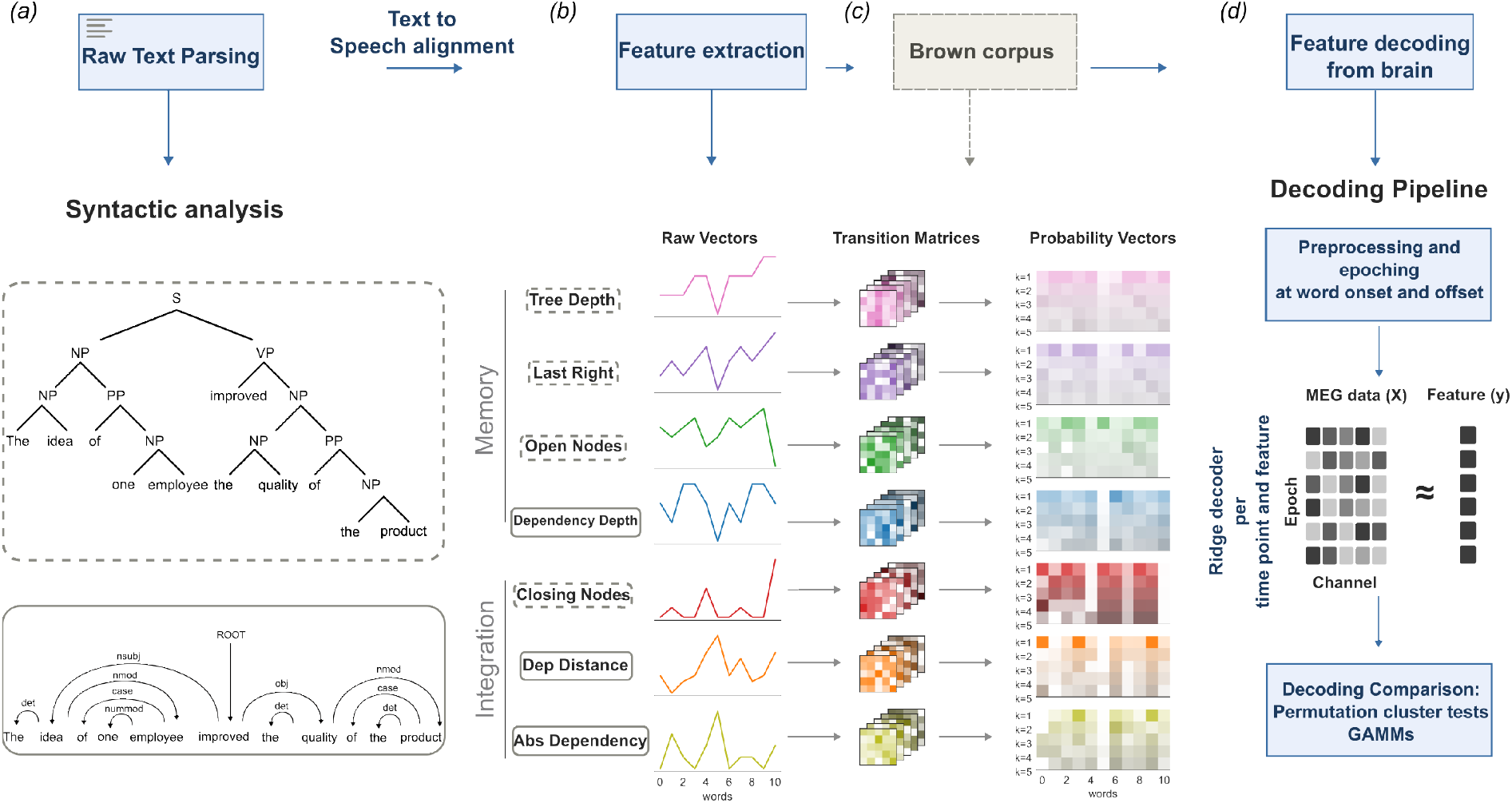
Decoding pipeline for memory and integration features. A. The raw text was parsed using both a constituency parser and a dependency parser to extract syntactic structures. B. Decoding features were extracted from syntactic structures by applying custom functions, and were grouped into memory and integration measures. Memory measures: *tree depth* quantifies how many nodes are above each word in the syntactic tree until the highest node is reached; *last right* adds +1 to tree depth whenever the word is the last syntactic item in a node (last right branch); *open nodes* quantifies how many nodes are still open for each word, subtracting the number of closing nodes; *dependency depth* quantifies the longest path between each word and the root of the sentence, i.e. the depth of the dependency tree. Integration measures: *closing nodes* counts how many nodes are resolved at each word; *dependency distance* calculates the distance between a dependent word and its head/governor, retaining the positive or negative sign that indicates dependency direction; absolute dependency distance is the same measure without a sign, and therefore without a direction. Feature extraction was applied to both transcriptions of the MEG dataset and the Brown Corpus [13]. C. Marginalized transition matrices were computed from the corpus and projected back onto the audiobook transcriptions. D. MEG data was minimally preprocessed and epoched at both word onset and offset. Each feature (both raw vectors and probabilities) was decoded independently, per time point. Finally, decoding scores were compared using permutation cluster tests and generalized additive mixed models.

The decodability of memory measures depended on the temporal anchor used for alignment (all p-values significant after Benjamini–Hochberg false-discovery-rate (FDR) correction). At word onset, *tree depth* (significant time points: 0.14 to 0.6 s, average t values in the cluster, 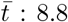, *p*_FDR_ *=0*.*0072* ) and *last right* features (0.14 to 0.6 s, 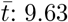, *p*_FDR_ *=0*.*0072* ) were successfully decoded. *Open nodes* was decoded within a smaller time window (0.16 to 0.31 s, 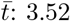, *p*_FDR_ *=0*.*0072* ). *Dependency depth* was decoded at a later time window (0.33 to 0.6 s, 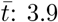, *p*_FDR_ *=0*.*0072* ) (see Fig. 2). At offset, only two memory measures were successfully decoded, but within later and smaller time windows: *tree depth* (0.34 to 0.6 s, 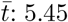, *p*_FDR_ *=0*.*0068* ); *last right* (0.23 to 0.6 s, 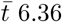, *p*_FDR_ *=0*.*0068* ). *Open nodes* was not decoded and the decoding performance of *dependency depth* was significantly below 0 (0.28 to 0.56 s, 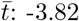, *p = 0*.*0068* ), suggesting a reversal of the discriminative pattern (e.g., high vs low dependency depth). Integration measures were similarly decoded at onset and offset: At onset, neural decoding was successful for *closing nodes* (0.13 to 0.6 s, 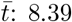, *p*_FDR_ *=0*.*0072* ), *dependency distance* (0.13 to 0.6 s, 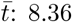, *p*_*FDR*_*= 0*.*0072* ), but not *absolute dependency* ; at offset, it was successful for all measures *closing nodes* (-0.03 to 0.6 s, 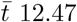, *p*_FDR_ *=0*.*0068* ), *dependency distance* (-0.04 to 0.6 s, 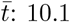, *p*_FDR_ *=0*.*0068* ), and *absolute dependency* (0.18 to 0.6 s, 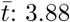, *p*_FDR_ *=0*.*0068* ).

**Fig. 2.**
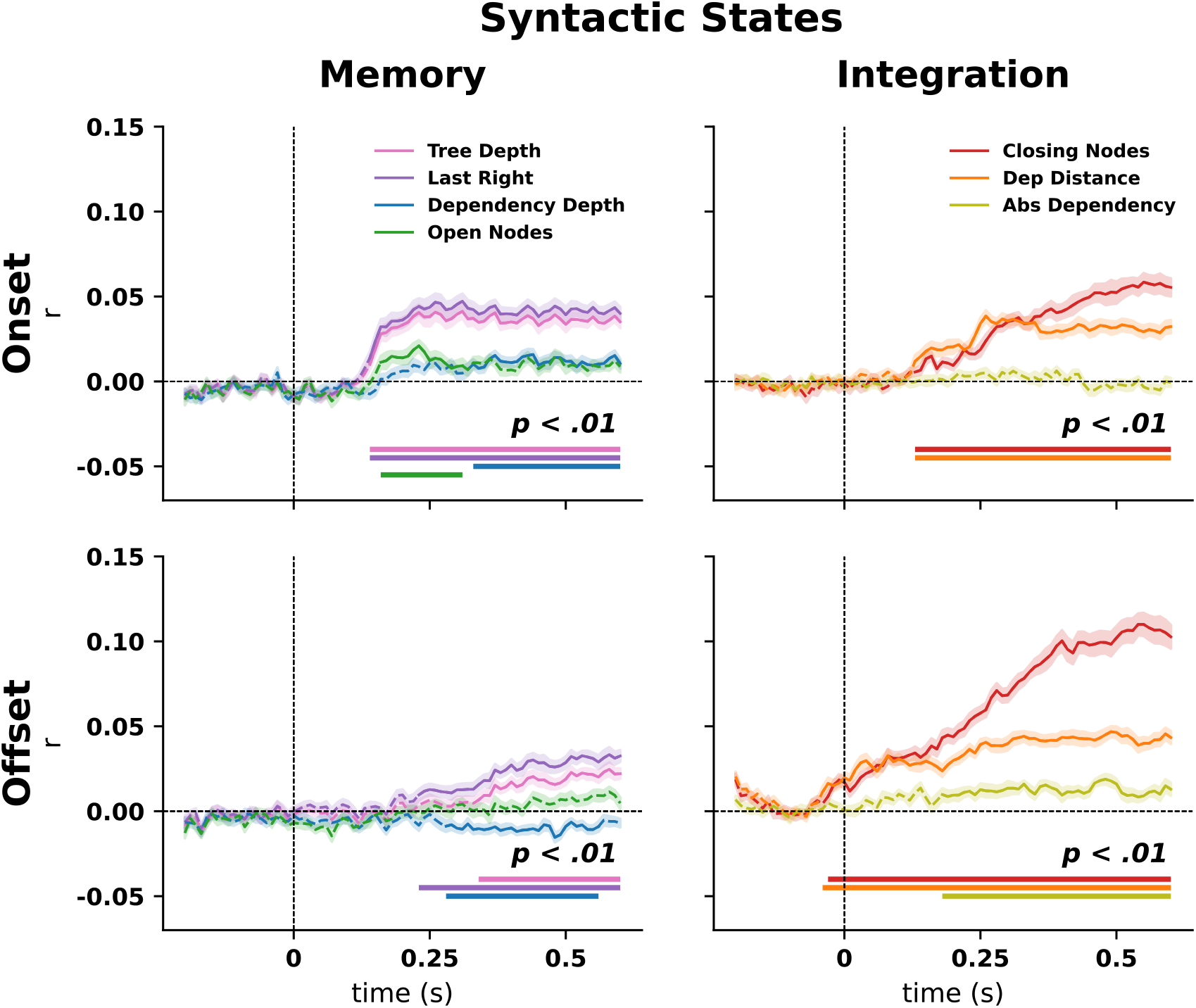
Decoding memory-based and integration-based syntactic structure at onset and offset. Raw features are individually color-coded. Solid lines indicate significance. Significance was computed against zero with a permutation cluster test. We report p values (after FDR correction) *<* .01.

### 2.2 Decoding syntactic expectations

To test the effects of prior knowledge, we modeled syntactic predictability by computing marginalized transition probabilities of syntactic weights using a Markov chain of first order for 5 lags or contexts (see Methods and Fig. 1 C), based on the Brown corpus [13]. We focused on *expectations*, so we chose to use *transition probabilities* instead of *surprisal*, which reflects prediction error [2, 3]. Using marginalized probabilities, we conditioned each estimate on one preceding item at a time rather than on the five contexts taken together, in order to determine their individual contribution to the distribution of decoding performance (Fig. 1 D). Projecting these probabilities onto the MEG dataset revealed that for memory-related syntactic measures, including *tree depth, last right, dependency depth* and *open nodes*, incorporating short-term contextual lags significantly improved decoding performance relative to base-line. Specifically, we found that both context -1 and context -2 enhanced decoding performance for all memory measures at onset (*tree depth* -1, -0.02 to 0.6 seconds 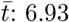, *p*_FDR_ *=0*.*006* ; *tree depth* -2, -0.02 to 0.6 s, 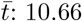, *p*_FDR_ *=0*.*006* ; *last right* -1,-0.04 to 0.6 s, 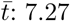, *p*_FDR_ *=0*.*006* ; *last right* -2, -0.02 to 0.6 s, 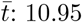 *p*_FDR_ *=0*.*006* ; *dependency depth -1*, 0.1 to 0.6 s, 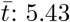, *p*_FDR_ *=0*.*006* ; *dependency depth -2* ; 0.1 to 0.6 s 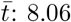, *p*_FDR_ *=0*.*006* ; *open nodes -1*, 0.25 to 0.6 s, 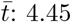, *p*_FDR_ *=0*.*006* ); *open nodes -2* -0.05 to 0.6 s, 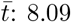, *p*_FDR_ *=0*.*006* ). Such performance enhancement was also observable at offset (*tree depth* -1, -0.06 to 0.52 s, 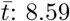, *p*_FDR_ *=0*.*007* ; *tree depth* -2, -0.06 to 0.6 s, 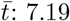, *p*_FDR_ *=0*.*007* ; *last right* -1, -0.05 to 0.39 s, 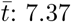, *p*_FDR_ *=0*.*007* ; *last right* -2, -0.06 to 0.45 s, 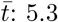, *p*_FDR_ *=0*.*007* ; *dependency depth* -1, -0.05 to 0.6 s, 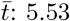, *p*_FDR_ *=0*.*007* ; *dependency depth* -2, 0.26 to 0.6 s, 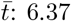, *p*_FDR_ *=0*.*007* ); *open nodes -1*, -0.04 to 0.6 s, 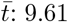, *p*_FDR_ *=0*.*007* ; context -2, -0.04 to 0.6, 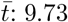, *p*_FDR_ *=0*.*007* ) (see Supplementary Materials B).

The effect of prior knowledge was not simply a sustained increase across all context, rather, it followed a non-monotonic pattern, highlighting a sharpening of locally predicted syntactic representations. Incorporating longer temporal contexts (contexts *-*4 and *-*5) reduced decoding performance to baseline values or even below baseline (e.g., at onset *tree depth* -4, 0.14 to 0.6 s, 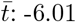, *p*_FDR_ *=0*.*006* ; *tree depth* -5, 0.11 to 0.6 s, 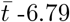, *p*_FDR_ *=0*.*006* ; *open nodes* - 4, 0.16 to 0.28 s, 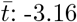, *p*_FDR_ *=0*.*006* ; *open nodes* -5, 0.14 to 0.44 s, 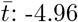, *p*_FDR_ *=0*.*006* ) (see Fig.3).

**Fig. 3.**
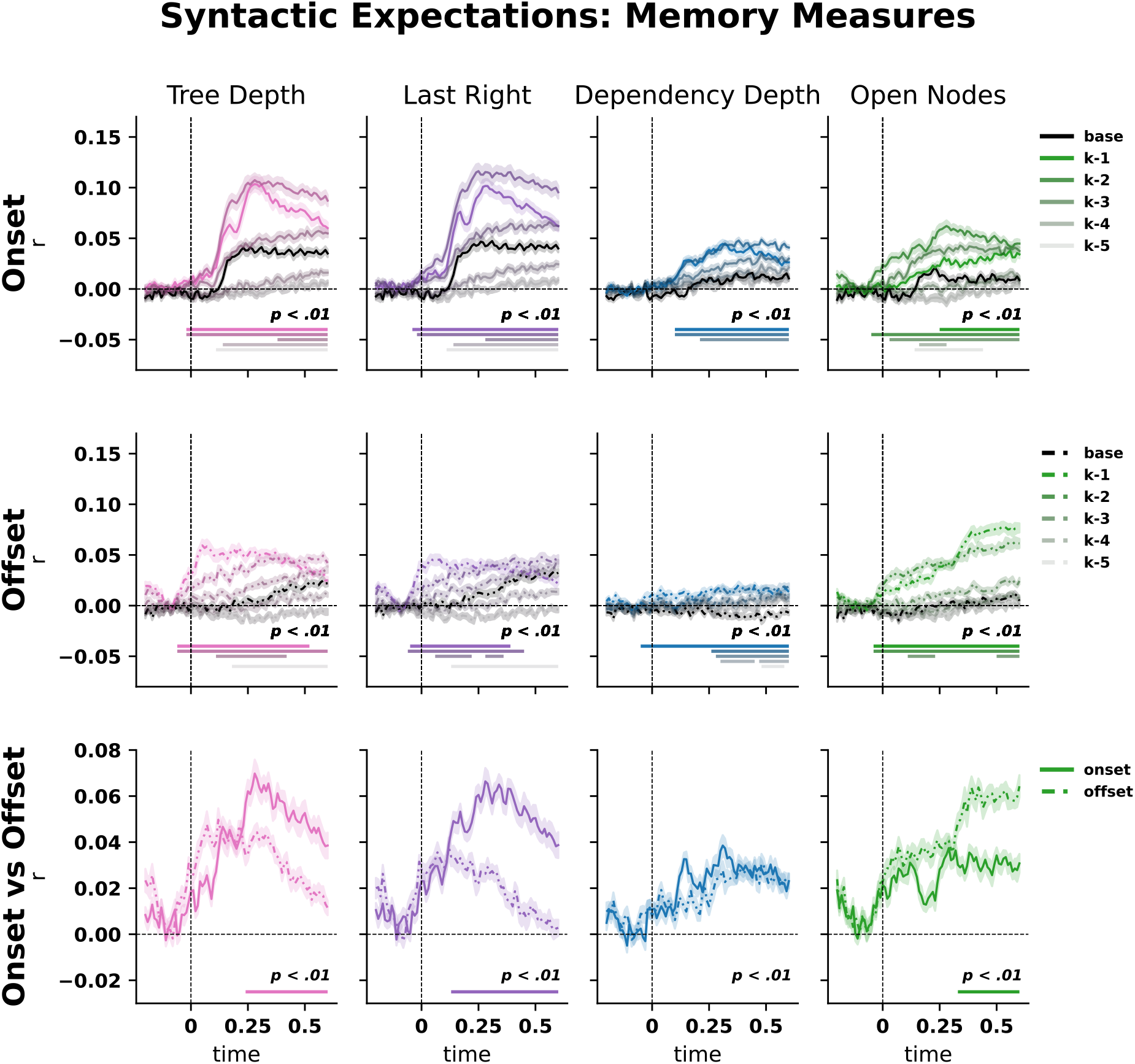
Sharpened memory-based syntactic structure via prior knowledge. Black lines indicate raw feature decoding performance. The significance of color-coded probability measures was computed against baseline vector decoding using a permutation cluster test, and was FDR corrected (*p*_*FDR*_ *<* .01). Color shading indexes the different contexts, while solid and dashed lines refer to onset and offset respectively. The last row shows the difference between onset and offset by computing the average of contexts (*k* ) -1 and -2, from which we first subtracted their respective baseline.

Such probabilistic sharpening of syntactic representations was feature-selective: integration-based measures failed to benefit from prior knowledge (see Fig. 4). Indeed, decoding performance for closing nodes and dependency distance remained consistently below baseline across almost all contexts, both at word onset and offset (e.g., at onset, *closing nodes -1*, 0.28 to 0.6 s, 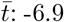, *p*_FDR_ *=0*.*006* ; context -2, 0.23 to 0.6 s, 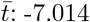, *p*_FDR_ *=0*.*006, dependency distance* -1, 0.25 to 0.6 s, 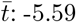, *p*_FDR_ *=0*.*006* ; context -2 0.24 to 0.6 s, 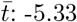, *p*_FDR_ *=0*.*006* ; see Supplementary Materials B). This dissociation provides neural evidence for the theoretical distinction between memory- and integration-based measures. Local syntactic expectations support the neural representation of syntactic features that await completion, but do not benefit those that complete a syntactic tree or one of its branches. We also run the same analysis using surprisal-based instead of probability-based contexts as further validation, replicating the dissociation between memory and integration measures with respect to their sensitivity to prior knowledge (see Supplementary Materials).

**Fig. 4.**
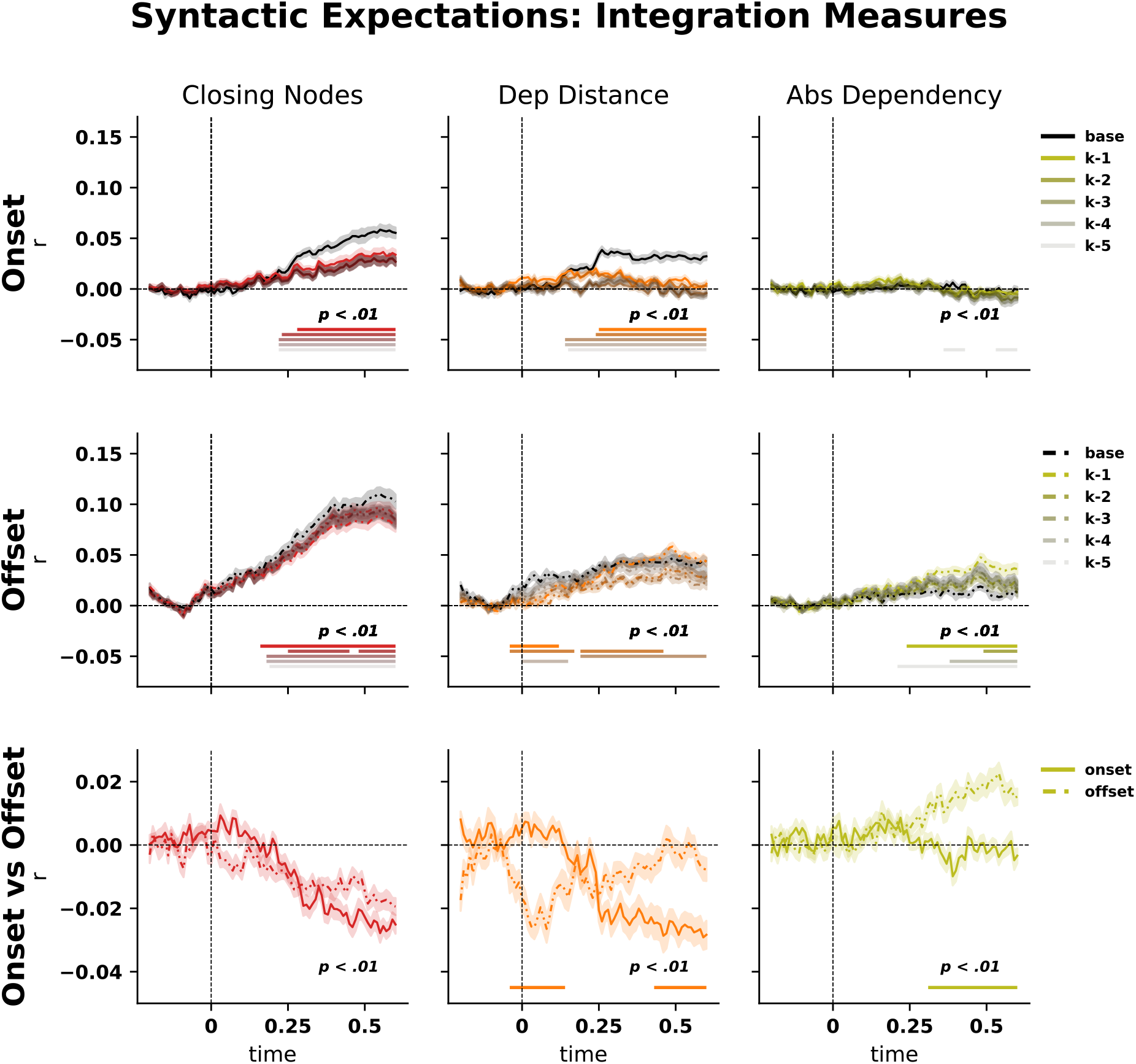
Prior knowledge effects on the neural decoding of integration-based syntactic information. Black lines indicate raw feature decoding performance. The significance of color-coded probability measures was computed against baseline vector decoding with a permutation cluster test, and FDR correction (*p*_*FDR*_ *<* .01). Color shading refers to the different contexts, while solid and dashed lines refer to onset and offset respectively. The last row shows the difference between onset and offset by computing the average of contexts (*k* ) *-*1 and *-*2, from which we subtracted their respective baseline.

### 2.3 Generalised sharpening

The cluster-based permutation analyses established that only nearby syntactic contexts selectively enhance the decoding of memory-related features. We asked whether the pattern we saw in individual contrasts reflects a broader, systematic property of the data, rather than a set of time-resolved paired contrasts. To address this, we analyzed the interaction between Expectation and Decoding feature factors separately for onset and offset using GAMMs. For memory features, the full GAMM provided a better fit than the reduced model with a shared smooth of time: onset, Δ *x*^2^ = 8.559, Δ df = 146.71, *p* < 2.2e-16); offset, Δ *x*^2^ = 3.098, Δ df = 100.31, *p <* 2.2e-16. Follow-up contrasts showed that nearby contexts were associated with enhanced decoding relative to baseline, whereas more distant contexts showed reduced decoding, consistent with a graded locality effect. This pattern was observed across memory-related features, although the timing and transition from facilitation to suppression varied across conditions (for a full table of contrasts, see Supplementary Materials D). Across models, the AR1 correction substantially reduced lag-1 residual autocorrelation, from *r* = [0.649] without correction to *r* = [0.032] after correction. Quadratic mixed-effects analyses of subject-level means confirmed this non-monotonic profile. Specifically, *dependency depth, last right, open nodes*, and *tree depth* all showed significant inverted U-shaped trends across baseline and probability contexts, both at onset and at offset (*p*_*FDR*_ *<* 0.01 in all cases, see Supplementary Materials E), with estimated turning points lying within the sampled context range and clustering around the early-to-intermediate probability levels (see Fig. 5). Thus, local sharpening is expressed as a systematic nonlinear property of the decoding trajectories and generalizes across memory-related features.

**Fig. 5.**
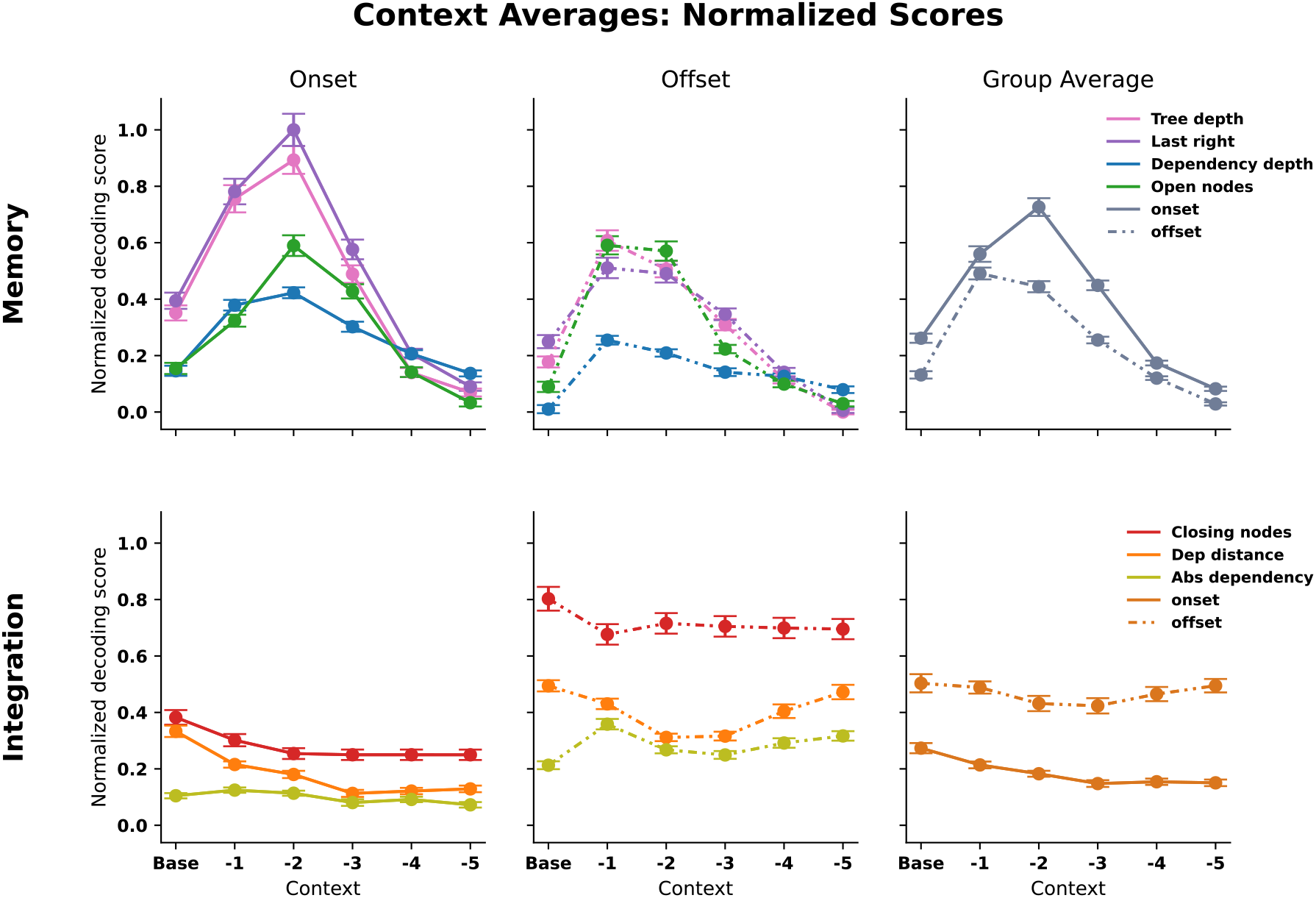
Normalized mean decoding scores for all measures and lags. Both at onset and offset, memory measures show a similar pattern of increase and decrease of decoding scores the follows an inverted U-shape response, indicating that information in the form of prediction is maintained in memory for a a brief time window before decaying. On the contrary, integration measures do not benefit from the prediction information, as the decoding scores remain stable across lags. Averaging all responses within group (memory and integration) also shows a dissociation between onset and offset, where memory and integration show opposite patterns. The figure was scaled using min-max scaling on the final mean decoding scores for the whole plate. Standard error was scaled by dividing SE by (max-min) values.

For integration measures, again the onset and offset analyses also favored the full model over the reduced model: onset, Δ *x*^2^ = 3.714, Δ df = 75.555, *p* < 2.2e-16); offset, Δ *x*^2^ = 4.791, 6df =101.26, *p* < 2.2e-16, indicating reliable context-dependent modulation of onset decoding. In contrast to memory measures, deviations from baseline were predominantly negative at onset, with the strongest effects observed for *dep distance* and *closing nodes*, and weaker effects for *abs dependency*, and heterogeneous across features at offset, with *abs dependency* and *closing nodes* predominantly positive relative to baseline, whereas *dep distance* showed a mixed pattern with negative deviations concentrated at intermediate contexts (for a full table of contrasts, see Supplementary Materials,D). Across models, the AR1 correction substantially reduced lag-1 residual autocorrelation, from *r* = [0.568] without correction to *r* = [0.036] after correction. Quadratic mixed-effects analyses did not provide evidence of an inverted U-shaped profile for any feature. In sum, locality effects differentiated memory and integration measures not only in sign, but also in their underlying trajectory shape.

To test whether onset- and offset-aligned decoding differed in their tendency to exhibit an inverted-U-shaped context profile across memory-related features, we fitted a quadratic mixed model to subject-level mean decoding values averaged across post-onset time points. The model revealed a significant phase × condition × quadratic-context interaction (*F* = 4.41, *p* = 0.0043), indicating that the difference in curvature between onset and offset depended on the decoded feature. Consistent with this result, the full model, which included phase-dependent linear and quadratic context interactions, provided a significantly better fit than the reduced model without these terms (Δ Deviance = 0.00847, Δ *df* = 7.00, *p* = 9.56 × 10^*-*13^). Follow-up feature-wise analyses showed negative quadratic coefficients at both onset and offset for all four memory-related measures. The phase-by-quadratic interaction survived false-discovery-rate correction for *dependency depth* (β = −0.00099, *p*_FDR_ = 0.0014), *last right (β* =−0.00291, *p*_FDR_ = 1.96 × 10^*-*6^), and *tree depth* (β = −0.00167, *p*_FDR_ = 0.0078), but not for *open nodes* (β = −0.00045, *p*_FDR_ = 0.386). In all significant cases, the quadratic coefficient was more negative at onset than at offset, indicating a stronger inverted-U-shaped context profile for onset-aligned decoding of memory-related syntactic features. Inspection of the phase-specific quadratic coefficients confirmed this pattern. For *dependency depth*, the quadratic coefficient changed from −0.00180 at offset to −0.00279 at onset; for *last right*, from −0.00366 to −0.00657; and for *tree depth*, from −0.00417 to −0.00583. By contrast, *open nodes* showed similarly negative curvature at offset and onset (−0.00440 and −0.00485, respectively), without a reliable onset*/*offset difference.

### 2.4 Anchor-dependent decoding

Finally, we asked whether differences between onset- and offset-based temporal anchoring reflected a transient effect confined to a limited temporal interval, or instead a more sustained property of neural decoding dynamics. To address this question, we compared decoding trajectories as a function of Alignment, Expectation, and Decoding feature factors, by fitting an omnibus subject × word anchor (alignment at onset or offset) × context (expectation modeled as marginalised probability bin) × condition (decoding feature) General Additive Mixed Model (GAMM [36]), separately for memory and integration feature sets. The omnibus model included an AR1 error structure to control for the inflating effects of residual autocorrelation across adjacent time points (see Methods). The full model - which allowed for onset- and offset-specific smooths within each context × condition combination - fit the data better than the reduced model, in which onset and offset shared the same temporal smooth: memory measures, Δ *x*^2^ = 5.796, Δ df = 134.18, *p* = 2.2e-16 ; integration measures, Δ *x*^2^ = 4.726, Δ df = 95.384, *p* = 2.2e-16. For memory features, positive onset-offset contrasts indicated generally stronger decoding at onset than at offset, with the largest differences observed for *last right* and *tree depth*, especially in context -1 and -2, followed by more moderate positive effects for *dependency depth. Open nodes* showed a weaker and less uniform pattern (see Figure 1, Supplementary Materials F). For integration features, negative onset-offset contrasts indicated stronger decoding at offset than at onset. The largest differences were observed for *closing nodes*, particularly in contexts -2 through to -5, followed by *dependency distance*, whereas *absolute dependency* showed the same pattern but with smaller effect sizes (see Figure 2, Supplementary Materials F). This pattern of results confirms and generalizes the findings obtained in decoding syntactic states: selecting a temporal anchor is a crucial step in language decoding, as it changes performance depending on feature type. Importantly, all onset–offset contrasts were robust after Benjamini–Hochberg false-discovery-rate (FDR) correction across context × condition cells (all *p*_*FDR*_ *<* 0.01, see Table 1 and Table 2 in Supplementary Materials F), indicating that onset/offset advantages generalized across the design space. We replicated the omnibus analysis for surprisal-based contexts, replicating the finding pattern for marginalized probabilities (see Supplementary Materials,G). Crucially, for both probability and surprisal measures, the onset-offset contrasts were typically sustained across large portions of the trajectory, rather than being confined to a specific temporal interval. This suggests that differences resulting from the temporal anchoring of neural decoding reflect an inherent property of neural processing. Furthermore, because the models included an AR1 error structure, it is unlikely that sustained differences merely reflected residual carry-over across adjacent samples, including silent intervals between words, rather than genuine differences in decoding dynamics.

**Table 1.**
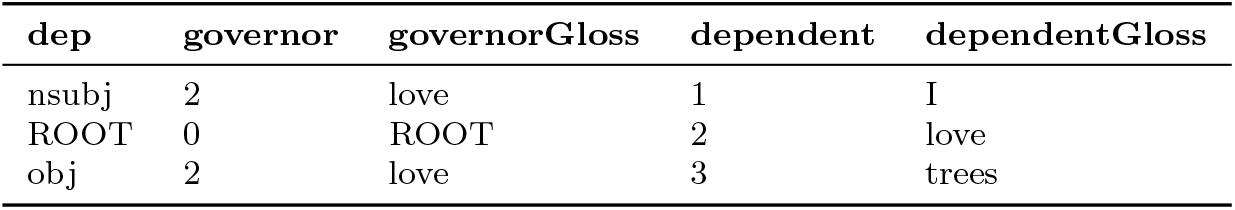
Dependency parse of the sentence “I love trees.”. Output from CoreNLP [50] API in [51].

## 3 Discussion

Understanding spoken language requires listeners to continuously infer syntactic structure from an unfolding and transient signal. Foundational neural work has shown that incremental comprehension depends not only on integrating each incoming word into an emerging representation, but also on generating probabilistic expectations about upcoming input[40, 41]. Here, we extend this account by showing that this process is not supported by a uniform gain in syntactic information. Prior syntactic knowledge selectively enhances the neural decoding of representations that must be maintained in working memory for upcoming input, whereas representations involved in the resolution of current structural relations show no comparable benefit from syntactic expectations. This dissociation suggests that syntactic structure building during online natural language comprehension is constrained by the computational role of the information being represented.

Our baseline decoding results - which index the current syntactic state - are consistent with previous evidence that abstract syntactic representations are encoded in neural activity during continuous language comprehension [9, 31, 33, 34]. We extend this work by decoding and combining features derived from both phrase-structure and dependency-based grammars, thereby moving beyond grammatical formalism to capture the functional profile of syntactic computations [30, 42]. In doing so, our approach helps bridge two largely separate traditions: NLP research, where dependency grammar has become a dominant formalism, and cognitive neuroscience, which has more often relied on phrase-structure accounts. More importantly, the ability to decode syntactic features across both grammatical traditions opens the way to asking not only whether syntax is represented in the brain, but how a competent listener operates over syntactic representations in real time. Competent listeners draw on prior knowledge about how linguistic structure is likely to unfold. To approximate these expectations, we estimated transition probabilities from a larger corpus. The effect was strikingly selective: prior knowledge enhanced the neural decoding of memory-related syntactic information, but not of information associated with structural resolution or integration, which was best captured by its raw, word-level code. This distinction is consistent with Dependency Locality Theory, which separates the cost of maintaining incomplete structure from the cost of integrating material once it becomes available. A core finding is that the enhancement of neural decoding for memory features was consistently local. Prior syntactic context improved decoding only over lags *-*1 and *-*2, while returning to baseline or worse over longer histories (see Fig. 5). Locality was preserved across word onset and offset, although the effect was stronger for alignment at word onset. We take this as evidence for a sharpening of the underlying syntactic representations, which are, by theory, locally bounded in computational scope. The resulting pattern, which was consistent across both probability-based and surprisal-based measures of syntactic expectation, suggests that in humans syntactic expectations are not maintained as a rich, long-range representation of the preceding sentence, but rather as a continuously updated, short-lived space of rule-based possibilities.

The strong, feature-selective locality effect in neural decoding links our findings to accounts that seek to unify prediction, memory and processing cost. In particular, lossy-memory accounts propose that the distinction between memory and integration costs may arise from degradation of representations over time [37, 43]. On this view, maintaining linguistic material is costly because past information becomes compressed, distorted or inaccessible as new input arrives. On the other hand, integrating linguistic material will have a higher cost at longer dependencies, modelled by the integration features. Our results are fully compatible with this framework: syntactic information that depends on maintenance shows a short-lived predictive benefit, whereas information associated with integration is only expressed when the relevant word is encountered. In sum, the data support a division between a forward-looking stream that maintains locally predicted structure, and a backward-looking operation set that resolves structure at the current syntactic state.

Our findings validate the use of audiobooks as representative samples for the study of human language complexity [44], and constrain a broad accounts of prediction in language comprehension. Prior knowledge is often treated as a general mechanism that optimizes the processing of upcoming information by pre-activating target neuronal ensembles. Our results suggest a more specific account: syntactic prediction enhances neural representations only when the predicted information must be maintained over time. Rather than simply work a gain function applied to all syntactic features, prior knowledge functions by stabilizing unresolved structure within the processing limits of the human language system.

The alignment-dependent temporal profiles of neural decoding further support this interpretation. Memory-related information was more strongly expressed using word onset as an anchor, consistent with a role for predictive structure in preparing the system for upcoming input. Integration-related information, by contrast, gained by anchoring to word offset, consistent with the idea that structural resolution unfolds once sufficient lexical and prosodic information has accumulated. These onset–offset differences should not be interpreted as a strict separation between prediction and integration in time. Rather, they reveal a graded shift in the dominant computational regime: early activity preferentially reflects maintained expectations, whereas later activity more strongly reflects the consequences of integrating the current word into the preceding structure.

Several limitations should be considered. First, decoding analyses of time-resolved MEG data can be inflated by temporal autocorrelation, because adjacent samples are not statistically independent. We addressed this issue by modeling the decoding trajectories with an AR1 error structure, reducing the risk that sustained effects reflect residual carry-over across neighboring time points. This is an important step for interpreting time-resolved decoding results, although future work could develop models that incorporate autocorrelation directly into the decoding stage, for example through time-series approaches such as ARIMA-based models [45]. Second, syntactic structure in natural speech is not independent of acoustic and prosodic structure. Phrase boundaries, pauses and pitch changes can all support syntactic segmentation, and integration-related measures may be especially correlated with silences at the ends of phrases or sentences. This could inflate absolute decoding performance for some integration measures, particularly around word offset. However, our main conclusions do not depend on decoding strength alone, but on the contrast between families of syntactic features and on their differential sensitivity to prior knowledge. Previous work also suggests that controlling for acoustic envelope responses does not necessarily eliminate syntactic decoding effects [9]. Still, the present results should not be taken to show that syntactic and prosodic information are fully separable in natural speech [35, 46]. Instead, they show that syntactic features with different computational roles exhibit distinct predictive and temporal profiles. Third, abstract linguistic features may covary with lower-level lexical properties, such as word length or frequency, which are also reflected in the MEG signal. We addressed this possibility via control analyses, which produced minimal changes in the observed pattern (see Supplementary Materials, I). Nevertheless, future work should model lexical, acoustic and syntactic structure jointly, especially in naturalistic designs where these dimensions are intrinsically correlated.

Finally, our findings have implications for the use of large language models (LLMs) as models of human language processing. LLMs’ predictions can align strongly with brain responses [8], but high alignment does not imply that humans and models rely on the same computational constraints. Recent evidence suggests that humans and LLMs differ in how they use contextual information, with humans compressing longer contexts into more limited representations [47]. Our results make this point for syntax, the core domain of structure building: when the target of the prediction is structured linguistic information, the relevant predictive context is strongly local (memory features), or even simply tributary to word-level operation (integration features). This helps explaining recent evidence that human predictions are bounded by syntactic domains, with stronger responses within phrases than across phrase boundaries [48]. In sum, the mechanism undelying neural language prediction in humans should not be equated with the mechanism responsible for maximal use of long textual context in LLMs. Human syntactic prediction appears powerful precisely because it is selective, compressed and constrained by the immediate demands of interpretation.

Together, these results show that the syntactic comprehension relies on a constrained predictive architecture: one that maintains what is locally useful, integrates what can be resolved, and discards or compresses what no longer serves the unfolding analysis.

## Methods

### 4.1 Parsing structures and dependencies

Arguably, two grammars have been the most popular in the field of NLP: phrase structure and dependency grammars. A phrase structure (or constituency) grammar consists of a formalization of combinatorics - that is, production rules - that specify how all syntactic items are related to each other [49]. These production rules are interpreted by a parser. The output of a phrase structure grammar is a nested structure represented as a graph tree where all terminal items are the words of the sentence, and each node carries a phrase label, which specifies the type of phrase. By design, a phrase node can contain one or more words/or one or more phrases.

Given the sentence:

1. I love trees A nested tree is conventionally represented in bracket notation as follows:
2. [S [NP [PRP I]] [VP [VBP love] [NP [NNS trees]]]]

In this notation, each node in the structure is marked by brackets. At each new open bracket, conventionally, the phrase label (S, NP, VP and so on) or the Part of Speech for words (PRP, VBP, NNS) is reported. The bracket notation corresponds to the tree structure reported in the tree example below. Dependency grammar is an alternative description of syntactic structure (see Table 1). This framework was originally introduced by [20] and became widely used in computational linguistics. Similarly to phrase structure grammars, dependency grammars formalize how words in a sentence related to each other. Specifically, two words (a governor and a dependent) are related by a dependency that characterizes what the relation formalizes (i.e., subject, object, case etc). Dependency grammars are still nested structures but can be represented as tables:

This specific aspect of the grammar substantially reduces the problem of dimensionality when vectorizing a sentence onto the time axis because there is no mismatch between the number of words and the number of words (as in a phrase structure grammar). This means that, in a scenario with continuous stimuli (both speech and text) the governor-dependent relation (marked by their word index) can be conceptualized as a sliding time window, for which the governor and dependent act as temporal delimiter.

Conventionally, both structure types are visualized as trees:

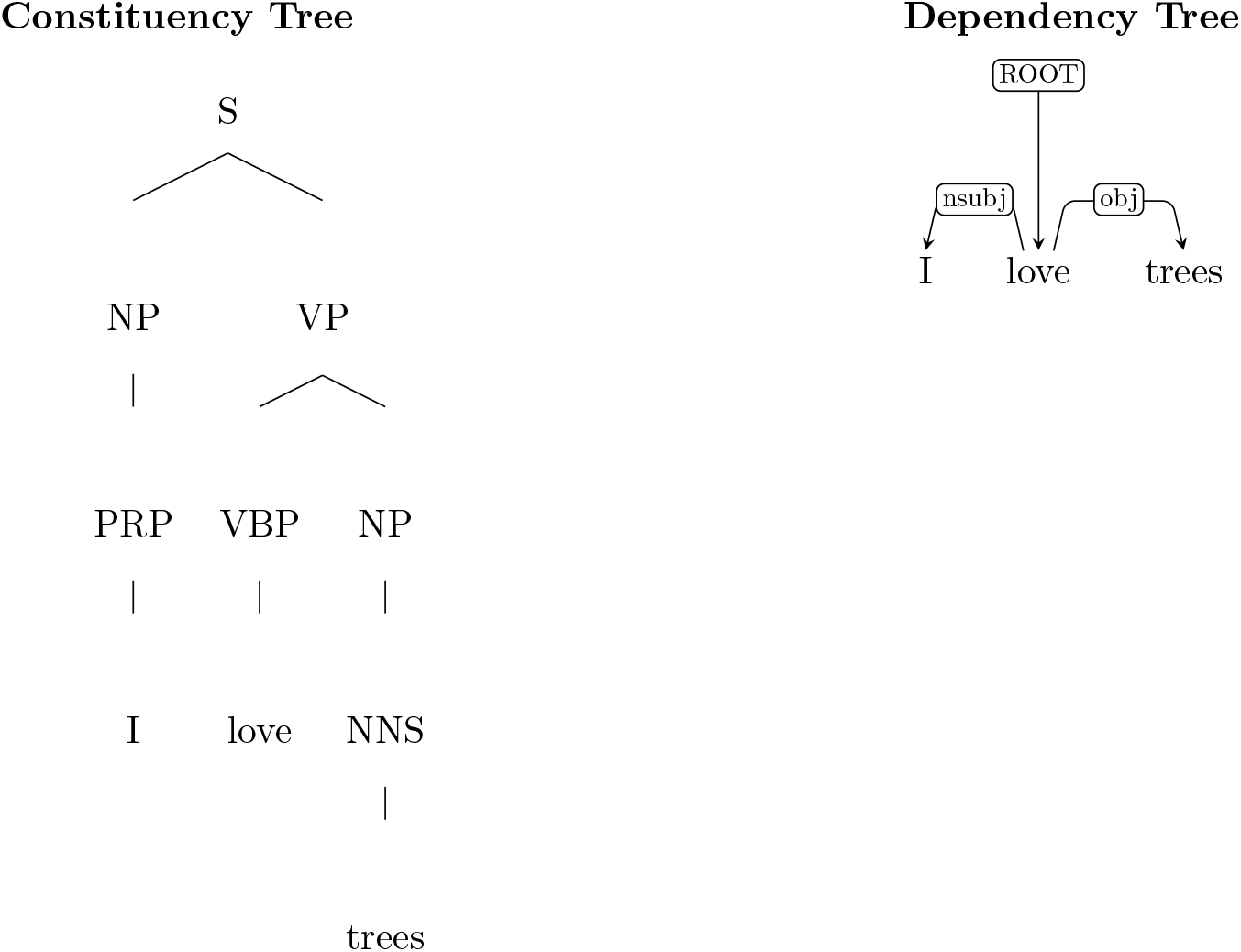

Comparing the two grammars, there are several differences both in terms of the information they capture and in terms of properties and limitations which have to be taken into account when using these formalism in cognitive neuroscience, paired with any time series (acoustics, EEG/MEG data, fMRI data). A phrase structure grammar describes mostly how words and phrases are grouped together but does not directly address the thematic/semantic relations among these words. A constituency tree does not directly show which words or phrases fulfill the role of subject (or object) in a sentence, although most of the times it can be inferred from the tree structure (an NP on the left and external to the main VP of the sentence is likely to be the subject). On the contrary, dependency grammars specifically address the relation among pairs of words in a sentence, by assigning a dependency label [52]. In both grammars, however, the word order will carry the temporal information (how words will unfold over time when speaking or listening) while additionally having a depth of the structure [16]. One final remark revolves around the number of nodes in both grammars: in a phrase structure grammar the number of nodes (i.e., NP, VP etc) can be different than the number of words, while in a dependency grammar each node is always represented by one word of the sentence (plus the ROOT). As a consequence, only dependency grammars can be truly described in a tabular form where each words receives a unique dependency label.

### 4.2 Defining a feature space through grammars

The study of sentence structure in the time domain (e.g., acoustics, M/EEG, or fMRI) requires the use of syntactically rich stimuli, such as sentences or full discourses. Especially in recent times, there has been a huge shift in experimental paradigms from very controlled stimuli (sentences that are matched for length and controlled for many dimensions) to long sessions where one can use a more ecologically valid setup [44]. By design, the use of naturalistic stimuli often requires the extraction of features to tag each unit in the stream, for example words. Mining such features from continuous stimuli, such as natural speech segments, is a non-trivial task as the design choices behind the extraction of linguistic information are usually tied to the theoretical background. This holds true for the study of any form of non physical properties of the speech signal, spanning from phonological processing (with the use of phonological features) to semantic processing, often addressed in recent literature with word embeddings coming from language models [9]. With this premise, the study of syntax also requires a representational space, i.e., any form of vectorization or feature space, to be time-locked to the acoustic signal or to the text, which can further be aligned to brain data. Indeed, by not having a contrast between highly controlled experimental conditions, it becomes crucial that we mark or characterize syntactic units with features that are theoretically informed. Syntactic structures are thought to be hierarchical in nature [53]. However, when presenting naturalistic stimuli to participants, speech unfolds linearly over the time axis. Most neuroimaging studies, especially using time sensitive techniques, will try to establish a statistical relation between two time series, an acoustic signal and a neural signal, in the form of, for example, a correlation between predicted and actual features (e.g., decoding literature, e.g. [9]) or mutual information between the acoustic envelope and the neural signal (speech tracking literature, e.g. [34]). In this framework, the first issue with modeling syntax concerns a problem of dimensionality [16, 17]: on one hand, abstract syntactic information is the result of nesting processes that will generate nested structures or trees, where the items (a lexical item or a phrase) will be bound together; on the other hand, the electro-physiological signal is subject to the constraint of time, unfolding on the temporal dimension. Regardless of the theoretical framework one chooses, both grammars generate structures which have a dimension that carries the word order and one dimension that carries the depth of structure. The first dimension, as it is the result of the linearization of a nested structure through time, can be matched to the time series of the acoustic signal, whereas the second dimension, which conveys the mapped relations among all units in the sentence, needs to be modeled to extract a vector of information. As a direct consequence, when studying sentence processing, the focus is on the structural dimension and how to quantify it over time (i.e., for every word). Here we provide a set of functions to replicate the extraction of measures of syntactic processing used in previous literature [9, 31, 38] and add the use of a dependency measure inspired by cross linguistic studies [18, 39]. We extend this automatic approach to more linguistic material to model how expectations on the upcoming structure during sentence processing can shape how nested structures are encoded by the brain. By comparing different approaches, we evaluate and quantify each measure of syntactic processing and whether it is relevant for the brain.

### 4.3 Design of the functions

We implemented a set of functions in Python, leveraging common NLP libraries, such as `stanza` [51], `NLTK` [54], and `CoreNLP` [50] client in `stanza` to parse the text. For the extraction of phrase structure measures (tree depth, last right, closing nodes, opening nodes) we transform the parse structures into strings to allow researchers to use different implementations, or to write their custom syntactic analysis, if needed. For the dependency measures, we use the CoreNLP client included in `stanza`. Additionally, the same client can be used in NLTK. Finally, we illustrate how dependency measures are computed, which can be replicated with other parsers and libraries once the dependency structure has been retrieved.

#### 4.3.1 Tree depth

In a constituency tree, the depth of the structure at each word corresponds to the number of nodes above that word until the highest node (usually S-node) is encountered. This corresponds to the longest path from the lexical item to the highest node in the structure. This measure describes how embedded each word is in the tree and can give a rough measure of how complex a structure is at each word point in the sentence. While being a proxy for the nesting process, the depth of the tree does not describe any *local dynamics* of the structure integration. For example, the tree structure in Fig 1A, shows that there is practically no difference among the words *The, idea*, and *of* in terms of depth, although the word *idea* actually collapses a Noun Phrase, and the word *of* belongs to a different phrase. Marking the depth of the structure of these three words results into all three having the same weight, despite the phrase boundary between *idea* and *of*.

#### 4.3.2 Last right (branch)

To account for local dynamics as well, we propose a measure that combines the depth of the structure and a minimal detection of phrase closing. Specifically, we use the depth of tree as a baseline and +1 whenever a word can be collapsed into a phrase, at any point in the structure. To compute these weights, we take the longest path from the root node to the current word and assume that if the word is in any position other than left, within a terminal node, it is closing a phrase. In some implementations of phrase structure grammars, a phrase can contain more than two elements (lexical items or other phrases). In such case, we add +1 for every non-left item in the phrase. This measure, by design, is highly correlated with the depth of the structure, and do not expect it to perform considerably better or differently than the depth of the structure. However, we still provide the implementation for future research.

#### 4.3.3 Closing nodes

The number of closing nodes at each word is one of the most used measures in combination with brain data. It was first introduced in neuroscience with fMRI data as a way to numerically describe the structure building over time [38] and later used in many studies [9, 33, 34, 55]. In a constituency tree, the number of closing nodes for each word is equal to the number of nodes between that word and the right-most terminal node is reached [38]. Going back to Fig 1A, the word *idea* closes one node, because the NP above it is the left branch of the higher NP. The word *employee*, instead, closes several nodes (NP, PP, and higher NP) because the first node being a left branch is the NP below the S node. As noted by Brennan et al. [38], counting the closing nodes corresponds to counting the number of closing brackets for each word in bracket notation. This measure has been proposed to be an indicator of how difficult is to process the sentence structure at each word-point.

#### 4.3.4 Open nodes

Opposite to closing nodes, the number of open nodes counts how many phrases have not been resolved at each word in a constituency tree. There have been several implementations of this measure [9, 31]. The number of open nodes for each word corresponds to the total number of nodes above that words from which we subtract the number of closed nodes. This measure quantifies how many nodes need to be maintained over the course of the sentence for each word.

#### 4.3.5 Dependency distance and Absolute dependency

In a dependency tree, each word is in a direct relation to another word, by means of a dependency. To measure the complexity of this dependency, we leveraged the approach in [39, 56, 57] and tagged every word with the number of words comprised between the dependent word and the head word. In such settings every word will receive a dependency but not every word will be the head of a dependency. As a consequence, some dependency go forward and some backward in the word order. More specifically given two words with index *j* and *i*, the dependency distance is quantified as the difference between the dependent index *j* and the head index *i*. Depending on how dependents and heads are ordered in the sentence, this will result into positive weights (the dependent precedes the head) or negative weights (the dependent follows the head). Additionally, we used the absolute dependency length (ignoring the sign of the dependency) as in [39].

#### 4.3.6 Dependency depth

Finally, we computed the depth of the dependency tree. As for the phrase structure grammar, the depth of the dependency tree corresponds to the longest path from one item to the highest node in the structure (the ROOT) [58]. Contrary to the phrase structure tree, each node in a dependency tree corresponds to one lexical item (excluding the ROOT). This means that the dependency depth count from word *i* to the root increases by one for each dependency needed to reach the highest node in the structure. Note that the dependency tree structure, as a result of having number of nodes equal to number of words (plus the ROOT), has generally a shallower structure than phrase structure trees.

### 4.4 Modelling sentence structure building through expectations

In order to define prior knowledge of syntactic structure of the English language, we implemented new measures of predictability based on the syntactic features we previously extracted. As a plausible representation of the English language, we took the Brown Corpus [13] contained in NLTK to scale our pipeline to a larger collection of text material to extract the raw syntactic features. The Brown Corpus is a collection of texts of different genres, ranging from novels to newspapers and beyond. It contained approximately 61000 sentences, for a total of ∼1.1 million words. We applied our feature extraction techniques to each sentence contained in the corpus separately, to extract all measures. Each of the 1.1 million words included in the corpus is now tagged with all the syntactic metrics proposed above (for the correlation structure of these features, see Supplementary Materials A). One of the criticism to the use of naturalistic stimuli concerns the genre of the stimulus material used during the experiment [44]. Particularly, it can be argued that audiobooks are representative only of a specific type of naturalistic stimuli and do not vary enough to actually represent natural language. This line of reasoning can be applied also for type of syntactic structures represented in audiobooks. As our pipeline is fully automatized, we can scale the extraction of these metrics beyond the single stories used in neuroscience experiment. To model the expectation of sentence structure building, we excluded punctuation from the feature space (around 10 % of all entries). First, we constructed transition probabilities of a specific syntactic weight given the preceding context. More precisely, we compute the marginalized transition probability of the syntactic weight *i* given *i-k*, with *k* ranging between 1 and 5, formalized as:

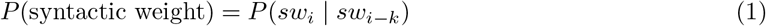

Note that this is a Markov chain of order one for each context because we are evaluating each word lag independently. Alternatively, one can compute transition probabilities conditioned on the full context. However, we decided against the latter option because i) we wanted to evaluate each word lag independently and ii) conditioning on full context might lead to sparser transition matrices. We then use the precomputed transition probabilities on the entire corpus and project it to the syntactic weight of the stories included in the MEG experiment and turn every metric in a measure of how likely is that the syntactic weight is indeed in that context within the sentence. These measures of predictability can be understood, at the conceptual level, as expectations of how the structure changes over time. Related to this, syntactic surprisal is often computed as the *-log*(*p*(*i*|*j*)), using as input to the transition matrix either the part of speech sequence [31], or the probability of a word given its context under the constraint of a phrase structure parser [1, 2]. In our approach, we chose to extract surprisal from the marginalized transition probabilities:

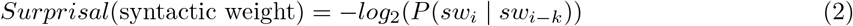

Deconding scores of syntactic surprisal are available in the Supplementary Materials C. In the main text we reported transition probabilities over surprisal as we are interested in *expectations* primarily. However, our full analysis revealed a very similar overall pattern using both transition probabilities and surprisal values.

### 4.5 Brain Dataset and Decoding

To validate our set of syntactic measures (both raw and probability measures) we performed a decoding analysis on an MEG dataset [59]. This dataset consists of 27 participants listening to 4 short stories in the format of audio. We excluded two participants as we were not able to import the data. The same stories were presented twice in 2 different sessions (not all participants took part in both sessions). To validate our feature space, we extracted the syntactic measures using the functions described above and aligned them to the word annotations included as metadata of the raw data: first, we parsed the transcriptions of the raw texts included with the dataset; second, we assigned transition probabilities at each word for different lags, based on the parsed raw text; third, we aligned the parsed structures to the words contained in the epochs. This step was necessary because the metadata of the MEG dataset miss a few words, as they were excluded from the original forced alignment procedure. To run the decoding analysis we adapted the code provided together with the dataset [59]. Data are minimally preprocessed as in the original paper, using mne [60]: first, we filtered the data between 0.5 and 30 Hz; then we constructed epochs at word onsets and offsets and down-sampled the data to 100 Hz. Each word is epoched at -0.2 seconds until 0.6 seconds after word onset and baseline correction is applied. Similarly, we epoched the data at word offset, computed as word duration added to word onset. The decoder is implemented as a Ridge Regression, which minimized the objective function:

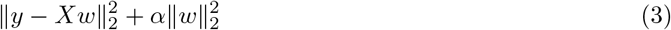

implemented in `sklearn` [61]. We kept alpha equal to 1, effectively applying a fixed penalty to all subjects and features, and allow for better comparability between features [62]. Features are scaled independently before fitting the decoder with `scale` and brain data were scaled in the cross validation loop using `Standard Scaler` from `sklearn`, respectively (z-score). Decoding was performed separately for each session, feature, and subject, concatenating all stories within session, using cross validation (K = 5). Decoding performance is evaluated by computing Pearson correlation per story and subject between the predicted features and the ground-truth features. Scores are averaged per story, session and subject, before a group average.

### 4.6 Statistical analysis: Cluster-based permutation tests

Statistical significance was evaluated using cluster-based permutation test. Raw features were tested using `permutation_cluster_1samp_test` included in `mne` [60], with default parameters, to test whether decoding was significantly better than chance. Cluster p-values were further corrected for multiple comparisons using the Benjamini–Hochberg false discovery rate approach (FDR). To evaluate whether transition probabilities or surprisal values lead to a significantly different decoding we used once again `permutation_cluster_1samp_tes`t for the decoding score of transition probabilities or surprisal from which we subtracted decoding scores of the raw features. We corrected once again for multiple comparisons by using the Benjamini–Hochberg FDR approach. Results are included only for corrected *p*-values *<* .01.

### 4.7 Statistical analysis: Generalized Additive Mixed Models

Decoding score trajectories were analysed using Generalized Additive Mixed Models (GAMMs) implemented with `bam()` in the `mgcv` package in R [36, 63]. The GAMM framework allows a flexible estimation of potentially nonlinear trajectories through penalised smooth functions, while accounting for repeated measurements across subjects.

We fitted separate GAMMs for onset- and offset-aligned decoding trajectories. For each phase, we first specified a reduced model in which decoding was allowed to differ across contexts and conditions in mean level, but shared a common temporal smooth:

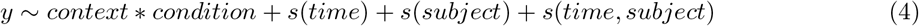

In `mgcv` notation, the reduced model (4) includes (i) a single smooth of time shared across all context × condition combinations, (ii) random intercepts for subjects (`s(subject, bs = “re”`)), and (iii) subject-specific smooth trajectories over time (`s(time, subject, bs = “fs”`)). We then compared this model to a full model, which allowed the temporal shape to vary separately for each context × condition combination:

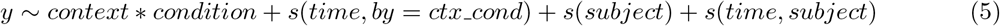

Here, `ctx_cond` denotes the interaction between context and condition. Thus, the full model (5) allows the temporal profile of decoding to vary across context Ccondition cells, rather than constraining all cells to share the same time course. Comparing the reduced and full models therefore tests whether contextual modulation is expressed in the shape of the decoding trajectory, rather than only in its overall level. To characterize the shape of the context effect at the subject level, we then computed mean decoding values for each participant within each context level, separately by feature and temporal anchor, averaging across all time points. These subject-level means were entered into quadratic mixed-effects models to test for non-monotonic context profiles. In these models, context was treated as an ordered predictor spanning baseline and the five probability levels, and a significantly negative quadratic term was taken as evidence for an inverted-U-shaped profile. Turning points were estimated from the fitted quadratic coefficients and evaluated with respect to the sampled context range. Formally, for each feature and temporal anchor, subject-level mean decoding values were modeled as:

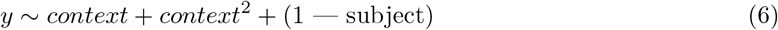

To test whether onset- and offset-aligned decoding differed in their tendency to generate an inverted-U-shaped context profile across memory features, we fitted an additional quadratic mixed-effects model to the same subject-level mean decoding values, including phase, condition, and quadratic context terms together with their interactions. We first specified a reduced model in which context exerted linear and quadratic effects that could vary across conditions, but were constrained to be the same for onset and offset:

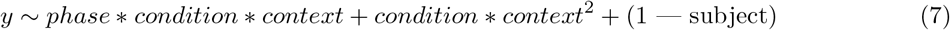

This model was compared against a full model that additionally allowed the quadratic effect of context to vary as a function of phase and condition:

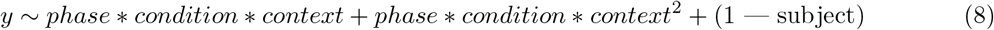

Comparing models (7) and (8) therefore tested whether onset- and offset-aligned decoding differed in the curvature of their context profiles as a function of decoded feature. Follow-up analyses then estimated phase-specific quadratic coefficients for each feature separately. For the integration features, the same GAMM and quadratic-modeling procedure was applied. This allowed us to assess whether context-related deviations from baseline followed the same nonlinear profile observed for memory features or instead expressed a distinct trajectory shape.

P values from follow-up feature-wise analyses were corrected for multiple comparisons using the Benjamini–Hochberg false-discovery-rate procedure, with significance assessed at *p*_*FDR*_ *<* 0.01. To test whether onset and offset trajectories differed in temporal shape as a function of context and condition, we fitted a model including fixed effects of *phase* (onset, offset), *context* (word position: base, -1:-5), *condition* (memory or integration measures), and their interaction, together with phase × context × condition-specific smooth functions of time, separately for memory and integration measures . We created a reduced model in which onset and offset were allowed to differ in mean level, but shared the same temporal smooth within each context × condition combination:

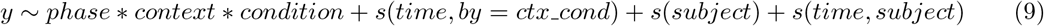

In `mgcv` notation, the reduced model (9) includes (i) separate smooths for each context × condition combination shared across onset and offset (`s(time, by = ctx cond`)), (ii) random intercepts for subjects (`s(subject, bs = “re”`)), and (iii) subject-specific smooth trajectories over time (`s(time, subject, bs = “fs”`)). The reduced model was compared to a full model, which allowed the temporal shape to vary separately for each phase × context × condition combination:

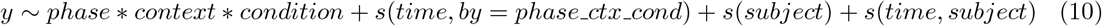

The factor `ctx_cond` denotes the interaction between context and condition, and `phase_ctx_cond` denotes the interaction between phase, context, and condition. Thus, the full model (10) allows onset and offset to differ not only in average level, but also in the shape of their time courses within each context × condition cell.

All models were fitted using restricted maximum likelihood (`fREML`). To reduce residual temporal dependence across adjacent samples within each subject × condition × context × phase trajectory, models included a first-order autoregressive error structure (AR1), specified through `rho` and `AR.start`. Full and reduced models were compared using approximate deviance-based *x*^2^ tests to account for residual auto-correlation across adjacent time points within each trajectory. This was important because decoding was analysed continuously over time, whereas the underlying linguistic signal was structured by word events which could be separated by brief silent intervals. Such temporal carry-over can induce short-range dependence between neighbouring samples and artificially inflate the apparent duration of decoding effects if left unmodelled. Models were fitted using `bam()` with a first-order autoregressive (AR1) error structure To further characterise temporal differences, for all models pairwise contrasts between smooths were computed from the model linear predictor matrix using `predict(…, type = “lpmatrix”`). These contrasts provided time-resolved differences, associated standard errors, and confidence intervals. For phase-specific condition-wise tests of context effects, model comparisons were performed separately within each condition, and resulting p-values were corrected for multiple comparisons using the false discovery rate (FDR) procedure.

## Acknowledgements

The authors would like to thank Benjamin Gagl and Bhavin Choksi for helpful comments on previous versions of the manuscript.

## Declarations

The authors declare no competing interests.

### Data availability

Raw MEG data are available on the Open Science Framework data repository at https://doi.org/10.17605/OSF.IO/AG3KJ. A description of data is available at [59].

### Code availability

Preprocessing and decoding scripts for the MEG data were readapted from https://github.com/kingjr/meg-masc/. All other relevant scripts are available at https://github.com/cosimo-iaia9305/Decoding-Syntactic-Expectations.

### Author contribution

Cosimo Iaia: Conceptualization, Methodology, Software, Validation, Formal Analysis, Writing - Original Draft; Alessandro Tavano: Conceptualization, Validation, Formal Analysis, Writing - Original Draft, Supervision.

### Source data

Source data for the figures are available at https://github.com/cosimo-iaia9305/Decoding-Syntactic-Expectations.

