## Supplementary Materials for "Prior knowledge reveals two computational regimes for syntactic processing in the human brain"

### Supplementary Information

#### Appendix A Descriptive statistics

In this section we report descriptive statistics of the stimulus set and features (Supplementary Fig. A 1, A2, and A3).

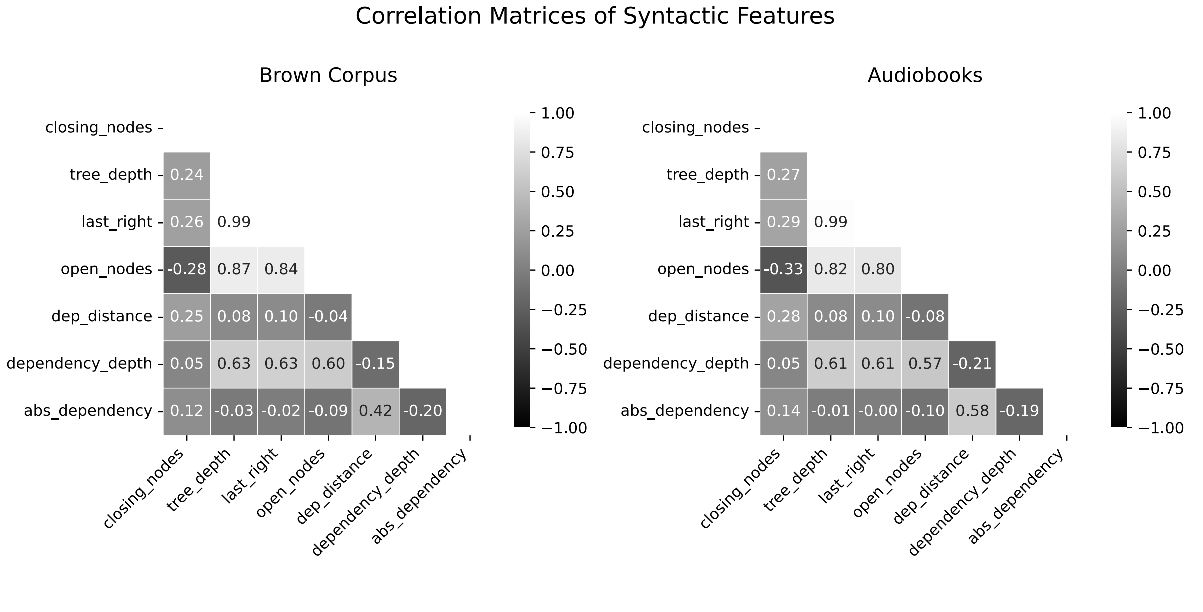


**Supplementary Fig. A 1** Correlation Matrices of syntactic features extracted both for the Brown Corpus and for the audiobooks contained in the MEG dataset.

#### Appendix B Decoding Transition Probabilities

Supplementary Fig. B 1 and B 2 report the distributions of transition probabilities. For each syntactic measure we generated 5 vectors, in which each element represent the transition probability of the original

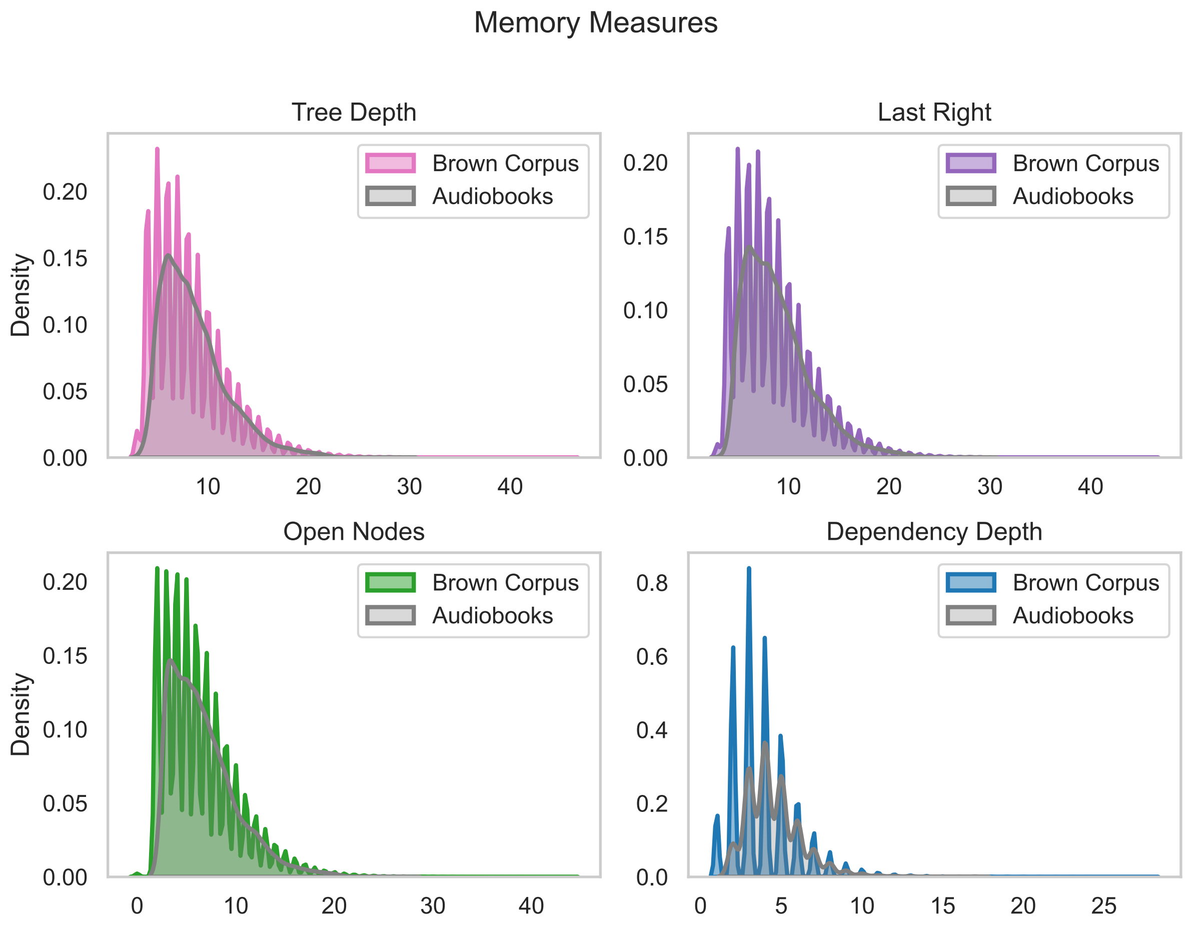


**Supplementary Fig. A 2** Density plot of memory measures. Colored lines describe the density plot of the Brown Corpus, while gray density plots show the distribution of the feature in the audiobooks of the MEG dataset. Gray distributions are offset for readability.

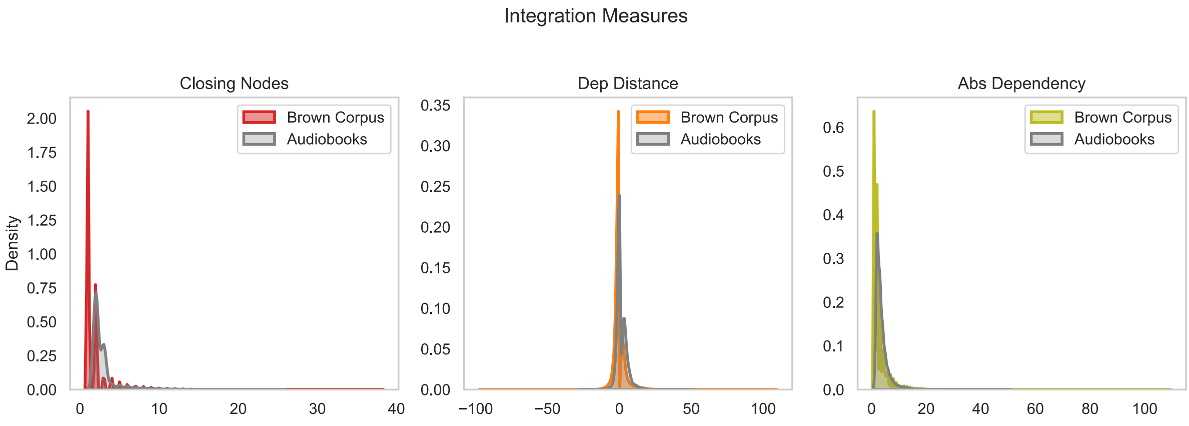


**Supplementary Fig. A 3** Density plot of integration measures. Colored lines describe the density plot of the Brown Corpus, while gray density plots show the distribution of the feature in the audiobooks of the MEG dataset. Gray distribution are offset for readability.

weight conditioned on the observed syntactic weight at position i-k. Results showed that incorporating transition probabilities improved decoding performance for memory-related syntactic measures (*tree* *depth*, *last right*, *dependency depth*, and *open nodes*) relative to baseline decoding, both at onset and offset. However, these improvements were not monotonic and were restricted to short temporal contexts. Specifically, performance enhancements were confined to contexts -1, -2, whereas longer contexts (-4 to -5) were associated with significant performance decreases. At onset, for both *tree depth* and *last right*, performance was significantly better than base decoding at context -1 and -2 (*tree depth* -1, -0.02 to 0.6 seconds $\bar{t}$: 6.93, $p_{\mathrm{FDR}}$ *=0.006*; *tree depth* -2, -0.02 to 0.6 s, $\bar{t}$: 10.66, $p_{\mathrm{FDR}}$ *=0.006*; *last right* -1,-0.04 to 0.6 s, $\bar{t}$: 7.27 , $p_{\mathrm{FDR}}$ *=0.006*; *last right* -2, -0.02 to 0.6 s, $\bar{t}$: 10.95 $p_{\mathrm{FDR}}$ *=0.006.* Similarly, context -3 shows a slight increase with a narrower cluster (*tree depth* - 3, 0.38 to 0.6 s, $\bar{t}$: 4, $p_{\mathrm{FDR}}$ *=0.006*; *last right* - 3, 0.28 to 0.6 s, $\bar{t}$: 4.6, $p_{\mathrm{FDR}}$ *=0.006*). Conversely, at earlier context, the same measure are significantly worse than base decoding (*tree depth* -4, 0.14 to 0.6 s, $\bar{t}$: -6.01, $p_{\mathrm{FDR}}$ *=0.006*; *tree depth* -5, 0.11 to 0.6 s, $\bar{t}$ -6.79, $p_{\mathrm{FDR}}$ *=0.006*; *last right* -4, 0.14 to 0.6 s, $\bar{t}$: -5.49, $p_{\mathrm{FDR}}$ *=0.006*; *last right* -5, 0.11 to 0.6 s, $\bar{t}$: -7.31, $p_{\mathrm{FDR}}$ *=0.006*). Similarly, at onset *dependency depth* is also significantly better than base decoding at context -1 (0.1 to 0.6 s, $\bar{t}$: 5.43, $p_{\mathrm{FDR}}$ *=0.006*), and at context -2 after (0.1 to 0.6 s $\bar{t}$: 8.06, $p_{\mathrm{FDR}}$ *=0.006*). At context -3, *dependency depth* is significantly better than base decoding (0.21 to 0.6 s, $\bar{t}$: 4.12, $p_{\mathrm{FDR}}$ *=0.006).* At context -4 and -5 *dependency depth* is not significantly different than base decoding. Lastly, open nodes at onset is significantly better at context -1 (0.25 to 0.6 s, $\bar{t}$: 4.45, $p_{\mathrm{FDR}}$ *=0.006*), at context -2 (-0.05 to 0.6 s, $\bar{t}$: 8.09, $p_{\mathrm{FDR}}$ *=0.006*) and at context -3 (0.03 to 0.6s $\bar{t}$: 6.35, $p_{\mathrm{FDR}}$ *=0.006*). At context -4 and -5, *open nodes* is significantly worse than base decoding (context - 4, 0.16 to 0.28 s, $\bar{t}$: -3.16, $p_{\mathrm{FDR}}$ *=0.006*; context -5, 0.14 to 0.44 s, $\bar{t}$: -4.96, $p_{\mathrm{FDR}}$ *=0.006*). For memory measures at the offset, we observed a similar sharpening, even though the base decoding performance was worse. At offset, for both *tree depth* and *last right*, performance was significantly better than base decoding at context -1 and -2 (*tree depth* -1, -0.06 to 0.52 s, $\bar{t}$: 8.59, $p_{\mathrm{FDR}}$ *=0.007*; *tree depth* -2, -0.06 to 0.6 s,$\bar{t}$: 7.19, $p_{\mathrm{FDR}}$ *=0.007*; *last right* -1, -0.05 to 0.39 s, $\bar{t}$: 7.37 , $p_{\mathrm{FDR}}$ *=0.007*; *last right* -2, -0.06 to 0.45 s, $\bar{t}$: 5.3, $p_{\mathrm{FDR}}$ *=0.007*). Context -3 also shows a increase in decoding performance (*tree depth* -3,0.11 to 0.42 s, $\bar{t}$: 3.71, $p_{\mathrm{FDR}}$ *=0.007*; *last right* -3, 0.06 to 0.22 s, $\bar{t}$: 3.09, $p_{\mathrm{FDR}}$ *=0.007*; 0.28 to 0.36 s, $\bar{t}$: 3.01, $p_{\mathrm{FDR}}$ *=0.007*). We did not observe an significant difference at context -4 for neither of these two measures at the offset. Conversely, at context -5, the same measure are significantly worse than base decoding (*tree depth* -5, 0.18 to 0.6 s, $\bar{t}$: -5.66, $p_{\mathrm{FDR}}$ *=0.007*; *last right* -5, 0.13 to 0.6 s, $\bar{t}$: -6.48, $p_{\mathrm{FDR}}$ *=0.007*). *Dependency depth* also benefits from a sharpening effect at all contexts (context -1, -0.05 to 0.6 s, $\bar{t}$: 5.53, $p_{\mathrm{FDR}}$ *=0.007*; context -2, 0.26 to 0.6 s, $\bar{t}$: 6.37, $p_{\mathrm{FDR}}$ *=0.007*; context -3, 0.28 to 0.6 s, $\bar{t}$: 4.53, $p_{\mathrm{FDR}}$ *=0.007*; context -4, 0.30 to 0.45 s, $\bar{t}$:3.97, $p_{\mathrm{FDR}}$ *=0.007*; context -4, 0.47 to 0.6 s, $\bar{t}$: 5.05, $p_{\mathrm{FDR}}$ *=0.007*; context -5, 0.48 to 0.58 s, $\bar{t}$: 3.73, $p_{\mathrm{FDR}}$ *=0.007*). Finally, *open nodes* shows a significant increase at word offset from modelling probabilities (context -1, -0.04 to 0.6 s, $\bar{t}$: 9.61, $p_{\mathrm{FDR}}$ *=0.007*; context -2, -0.04 to 0.6, $\bar{t}$: 9.73, $p_{\mathrm{FDR}}$ *=0.007*; context -3, 0.11 to 0.23, $\bar{t}$: 2.73, $p_{\mathrm{FDR}}$ *=0.007*; context -3, 0.5

to 0.6 s, $\bar{t}$: 4.13, $p_{\mathrm{FDR}}$ *=0.007*). In contrast, integration-based measures did not benefit from transition probabilities at onset and offset, indicating a selective effect of syntactic predictability. At onset, *closing nodes* is consistently lower than its base decoding (context -1, 0.28 to 0.6 s, $\bar{t}$: -6.9, $p_{\mathrm{FDR}}$ *=0.006*; context -2, 0.23 to 0.6 s, $\bar{t}$: -7.014, $p_{\mathrm{FDR}}$ *=0.006*, context -3, 0.22 to 0.6 s, $\bar{t}$: -7.36, $p_{\mathrm{FDR}}$ *=0.006*; context -4, 0.22 to 0.6 s, $\bar{t}$: -7.34, $p_{\mathrm{FDR}}$ *=0.006*; context -5, 0.22 to 0.6 s, $\bar{t}$: -7.37, $p_{\mathrm{FDR}}$ *=0.006*). Similarly, *dependency distance* becomes significantly lower than base decoding at each possible context window (context -1, 0.25 to 0.6 s, $\bar{t}$: -5.59, $p_{\mathrm{FDR}}$ *=0.006*; context -2 0.24 to 0.6 s, $\bar{t}$: -5.33, $p_{\mathrm{FDR}}$ *=0.006*, context -3, 0.14 to 0.6 s, $\bar{t}$: -6.65, $p_{\mathrm{FDR}}$ *=0.006*; context -4, 0.14 to 0.6 s, $\bar{t}$: -6.59, $p_{\mathrm{FDR}}$ *=0.006*; context -5, 0.15 to 0.6 s, $\bar{t}$: -6.91, $p_{\mathrm{FDR}}$ *=0.006*). Lastly, *absolute dependency* remains very poorly decodable even when looking at local contexts and becomes significantly worse at context -5 (0.36 to 0.43 s, $\bar{t}$: -3.03, $p_{\mathrm{FDR}}$ *=0.006*, 0.53 to 0.6 s, $\bar{t}$: -3.79, $p_{\mathrm{FDR}}$ *=0.006*). At offset, *closing nodes* is also consistently lower than its base decoding (context -1, 0.16 to 0.6 s, $\bar{t}$: -6.34, $p_{\mathrm{FDR}}$ *=0.007*; context -2, 0.25 to 0.45 s, $\bar{t}$: -3.65, $\mathrm{p}_{\mathrm{FDR}}$ =0.007; context -2, 0.48 to 0.6 s, $\bar{t}$: -4.44, $p_{\mathrm{FDR}}$ *=0.007*; context -3, 0.18 to 0.6 s, $\bar{t}$: -4.43, $p_{\mathrm{FDR}}$ *=0.007*; context -4, 0.18 to 0.6 s, $\bar{t}$: -4.71, $p_{\mathrm{FDR}}$ *=0.007*: context -5, 0.19 to 0.6 s, $\bar{t}$: -5.03, $p_{\mathrm{FDR}}$ *=0.007*). Similarly, *dependency distance* becomes significantly lower than base decoding at each possible context window (context -1, -0.04 to 0.12 s, $\bar{t}$: -4, $p_{\mathrm{FDR}}$ *=0.007*; context -2, -0.04 to 0.17 s, $\bar{t}$: -4.62, $p_{\mathrm{FDR}}$ *=0.007*; context -2, 0.19 to 0.46 s, $\bar{t}$: -4.68, $p_{\mathrm{FDR}}$ *=0.007*; context -3, 0.19 to 0.6 s, $\bar{t}$: -4.62, $p_{\mathrm{FDR}}$ *=0.007*; context -4, 0.0 to 0.15 s, $\bar{t}$: -3.13, $p_{\mathrm{FDR}}$ *=0.007*), while not being significantly different at context -5. Lastly, *absolute dependency* shows an improvement compared to baseline at offset for some contexts (context- 1, 0.24 to 0.6 s, $\bar{t}$: 6.01, $p_{\mathrm{FDR}}$ *=0.007*; context- 2, 0.49 to 0.6 s, $\bar{t}$: 3.29, $p_{\mathrm{FDR}}$ *=0.007*; context- 4, 0.38 to 0.6 s, $\bar{t}$: 4.34, $p_{\mathrm{FDR}}$ *=0.007*; context- 5, 0.21 to 0.6 s, $\bar{t}$: 4.77, $p_{\mathrm{FDR}}$ *=0.007*). Finally, we compared the enhancement effect between onset and offset by averaging within each measure the results of context -1 and context -2, from which we then subtracted the base decoding. We observed an increase in all memory measures, with some difference between onset and offset for three of them (*tree depth* enhancement onset vs offset, 0.24 to 0.6 s, $\bar{t}$: 4.37, $p_{\mathrm{FDR}}$ *=0.0073*; *last right*, 0.13 to 0.6 s, $\bar{t}$: 6.04, $p_{\mathrm{FDR}}$ *=0.0073*; *open nodes*, 0.33 to 0.6 s, $\bar{t}$: -5.43, $p_{FDR}$ *=0.007= 0.0073*). Conversely, for integration measures, we observe significant difference for *dependency distance* (offset significantly more decreased than onset, -0.04 to 0.14 s, $\bar{t}$: 4.15, $p_{\mathrm{FDR}}$ *=0.0073*; onset significantly more decreased than offset, 0.43 to 0.6 s, $\bar{t}$: -3.67, $p_{\mathrm{FDR}}$ *=0.0073*) while absolute *dependency distance* shows with an increase for offset compared to onset (0.31 to 0.6 s, $\bar{t}$: -4.55, $p_{\mathrm{FDR}}$ *=0.0073*). We observed no significant change for closing nodes.

### Appendix C Decoding Surprisal

To check if surprisal has a similar impact on the decoding scores, we ran an additional analysis where instead of decoding the transition probabilities of each syntactic weight, we decode their surprisal values

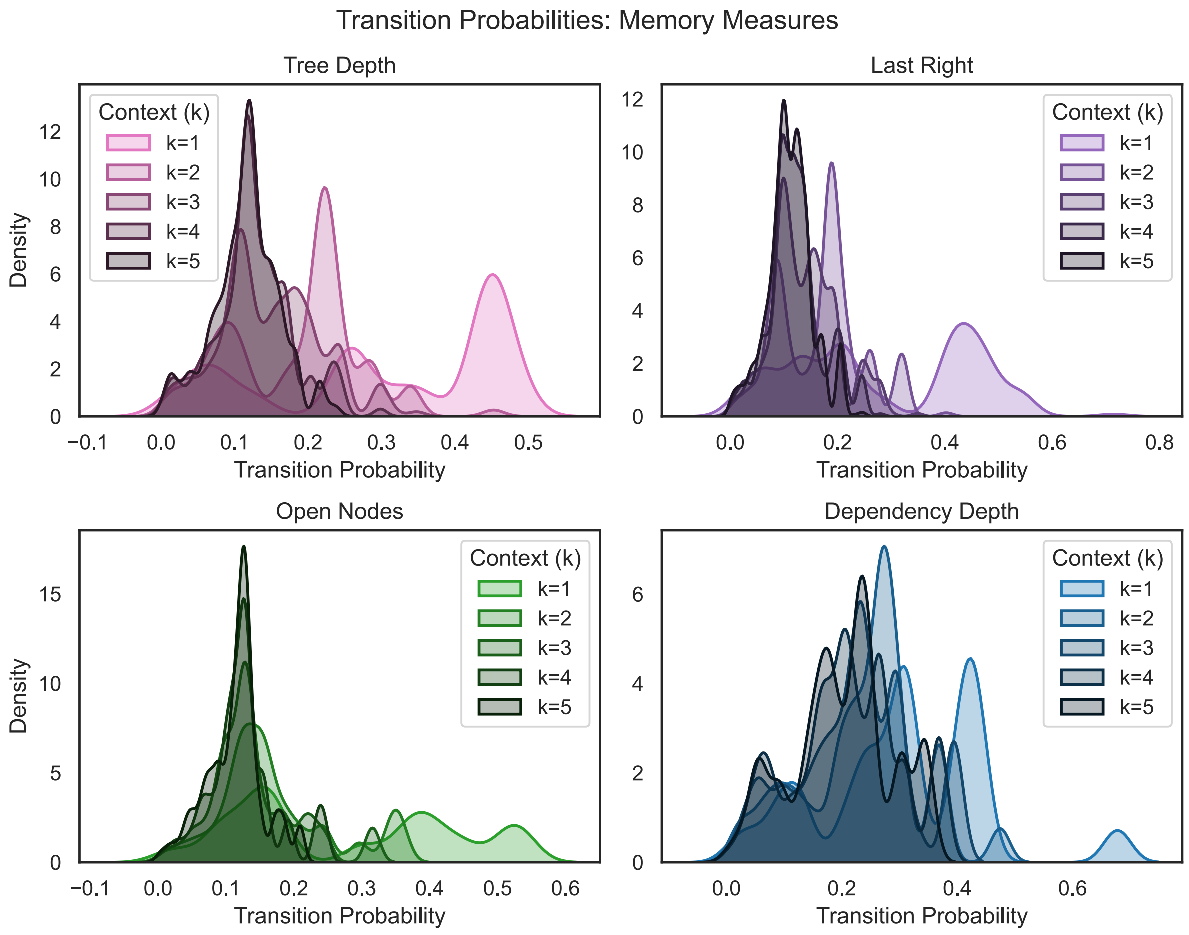


**Supplementary Fig. B 1** Distribution of transition probabilities of memory features ad different lags. K indicates the actual lag of the distribution.

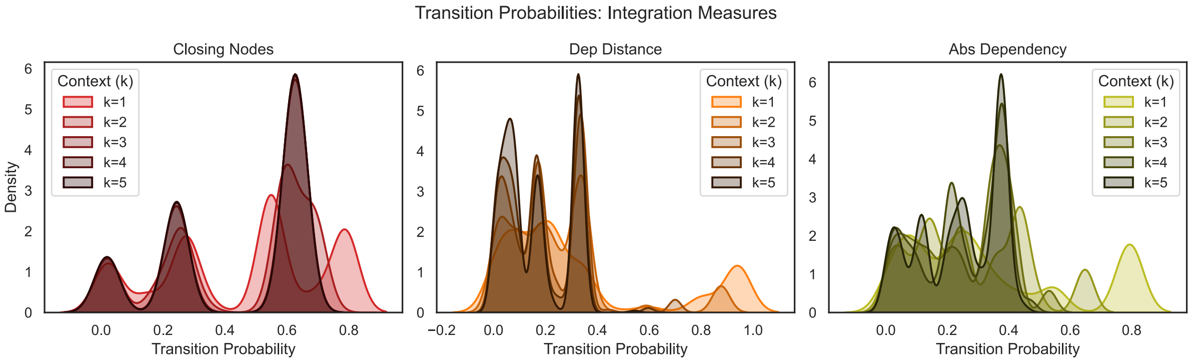


**Supplementary Fig. B 2** Distribution of transition probabilities of integration features ad different lags. K indicates the actual lag of the distribution.

Supplementary Fig. C1, C2, C3 and C4. We computed surprisal values by using $-\log 2(p(sw_i|sw_{i-k}))$. We repeated this analysis for both memory measures and integration measures, at onset and offset. At onset, we found similar results to using probability at context length 1, producing a sharpening of memory measures. *Tree depth* and *last right* are significantly higher than base decoding only at context -1 (0.03

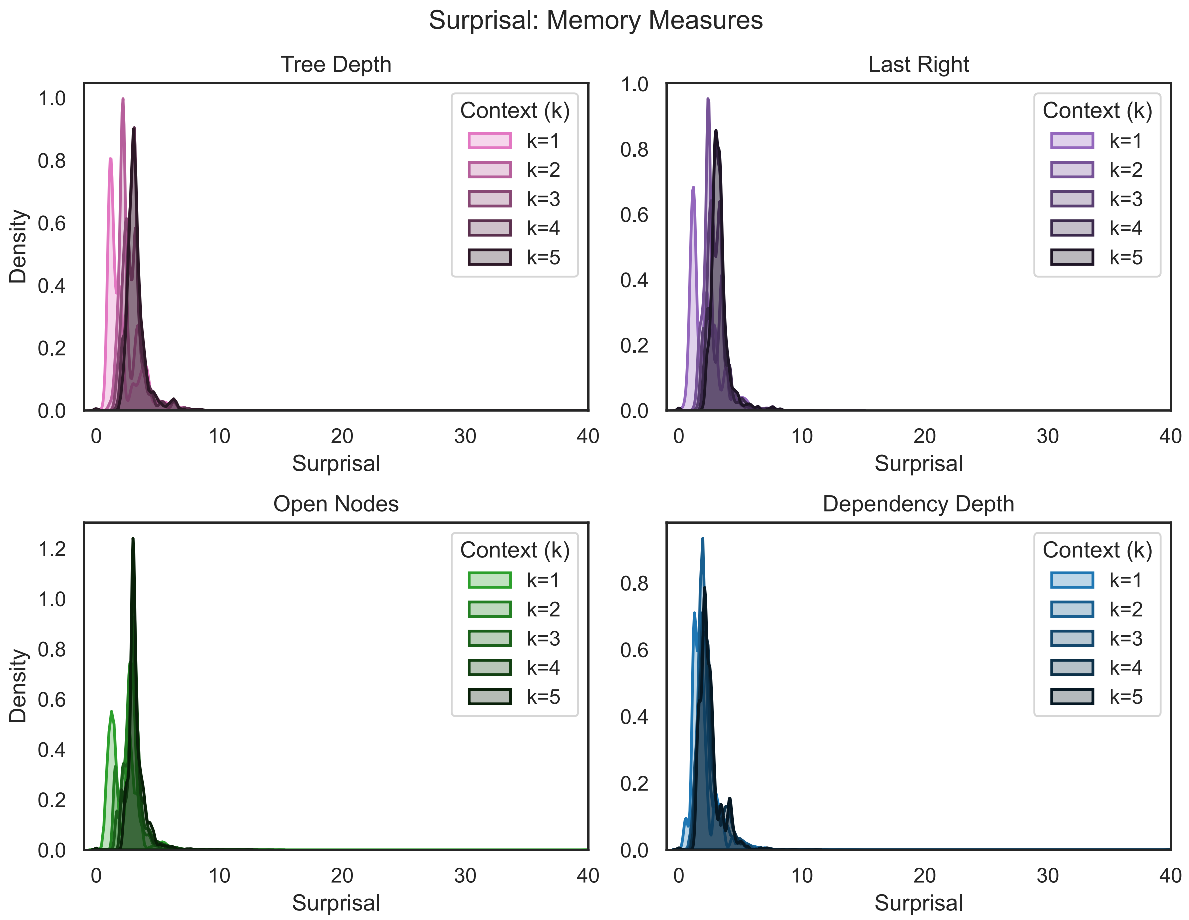


**Supplementary Fig. C 1** Density plot of surprisal values for expectation measures

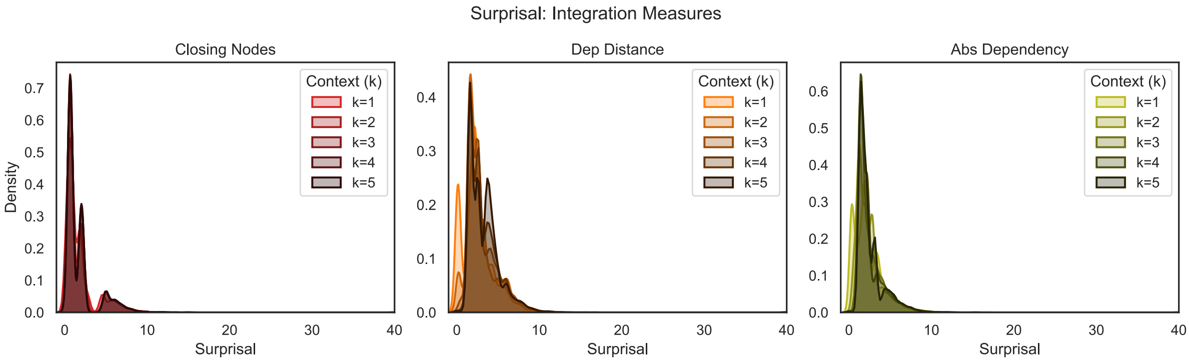


**Supplementary Fig. C 2** Density plot of surprisal values for integration measures.

to 0.6 s, $\bar{t}$: 9.07, $p_{\mathrm{FDR}}$ $=0.008$; 0.03 to 0.6 s $\bar{t}$: 9.23, respectively, $p_{\mathrm{FDR}}$ $=0.008$). Similarly, at context -2, both measures are significantly better than base decoding (*tree depth*, 0.1 to 0.6 s, $\bar{t}$: 8.34, $p_{\mathrm{FDR}}$ $=0.008$; *last right*, -0.01 to 0.6 s, $\bar{t}$: 8.36, $p_{\mathrm{FDR}}$ $=0.008$). In contrast, longer contexts become significantly worse at onset for *tree depth* (context -4, 0.13 to 0.6 s, $\bar{t}$: -6.51, $p_{\mathrm{FDR}}$ $=0.008$; context -5, 0.13 to 0.6 s, $\bar{t}$: -6.71, $p_{\mathrm{FDR}}$ $=0.008$) and *last right* (context -4, 0.12 to 0.6 s, $\bar{t}$: -7.67, $p_{\mathrm{FDR}}$ $=0.008$; context -5, 0.09 to 0.6

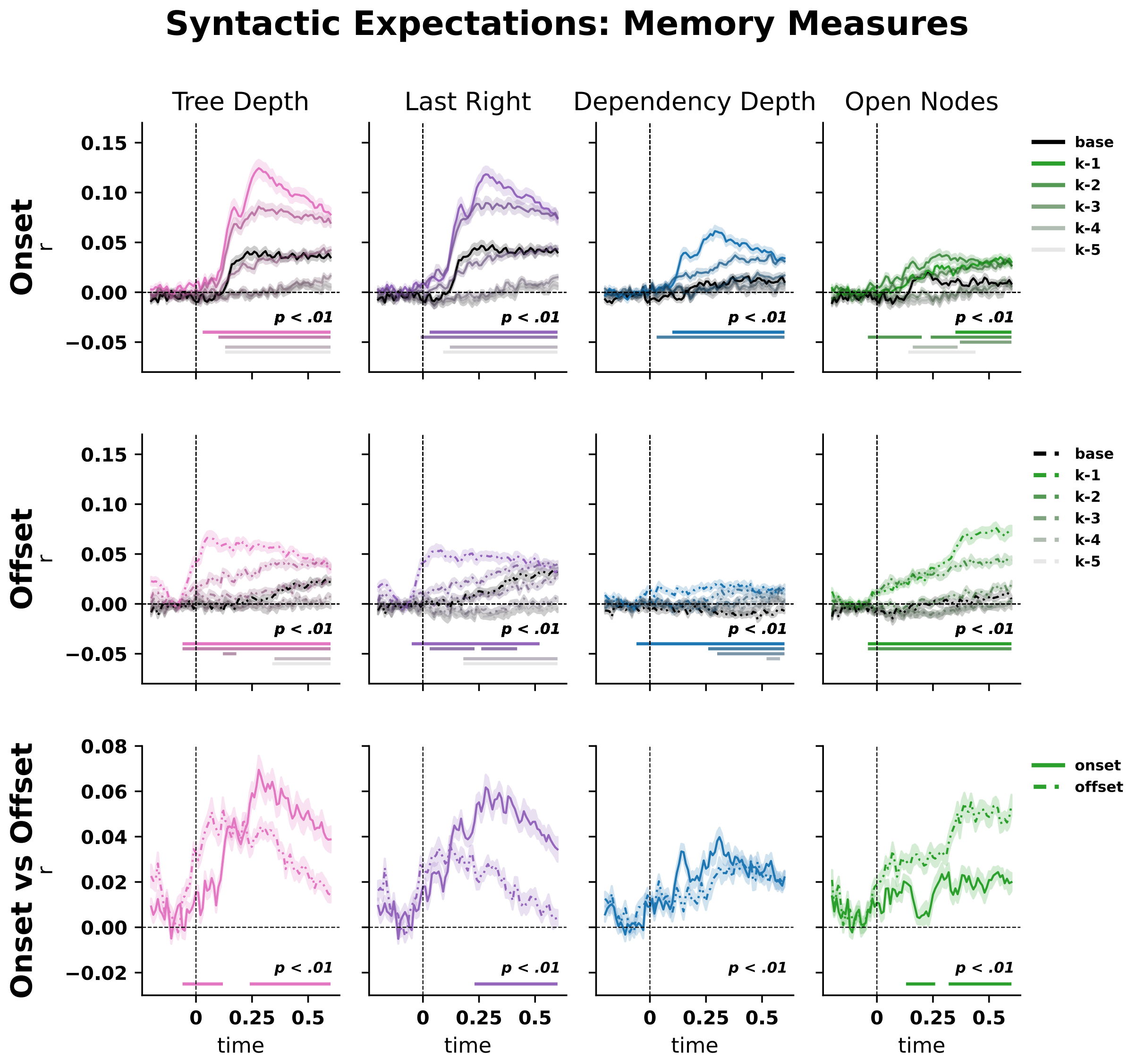


**Supplementary Fig. C 3** Sharpened memory-based syntactic structure using surprisal values. Black lines indicate raw syntactic state feature decoding performance. The significance of color-coded probability measures was computed against baseline vector decoding with a permutation cluster test, and FDR correction ($p_{FDR}$ <.01). Color shading refers to the different contexts, while solid and dashed lines refer to onset and offset respectively. The last row shows the difference between the context -1 and -2 against the base decoding for both onset and offset

s $\bar{t}$: -7.57, $p_{\text{FDR}}$ $=0.008$). At context -3, neither *tree depth* nor *last right* is significantly different than baseline.

Similarly, *dependency depth* at onset shows an enhancement of decoding performance at context -1 (0.1 to 0.6 s, $\bar{t}$: 7.28, $p_{\text{FDR}}$ $=0.008$) and context -2 (0.03 to 0.6 s, $\bar{t}$: 4.98, $p_{\text{FDR}}$ $=0.008$), while not changing significantly at contexts -3, -4, and -5. *Open nodes* is significantly better than base decoding, supporting a locality effect (context -1, 0.35 to 0.6 s, $\bar{t}$: 4.31, $p_{\text{FDR}}$ $=0.008$; context -2, -0.04 to 0.2, $\bar{t}$: 4.37, $p_{\text{FDR}}$ $=0.008$; context -2, 0.24 to 0.6 s, $\bar{t}$: 5.11, $p_{\text{FDR}}$ $=0.008$, context -3, 0.37 to 0.6s, $\bar{t}$: 4.31, $p_{\text{FDR}}$ $=0.008$),

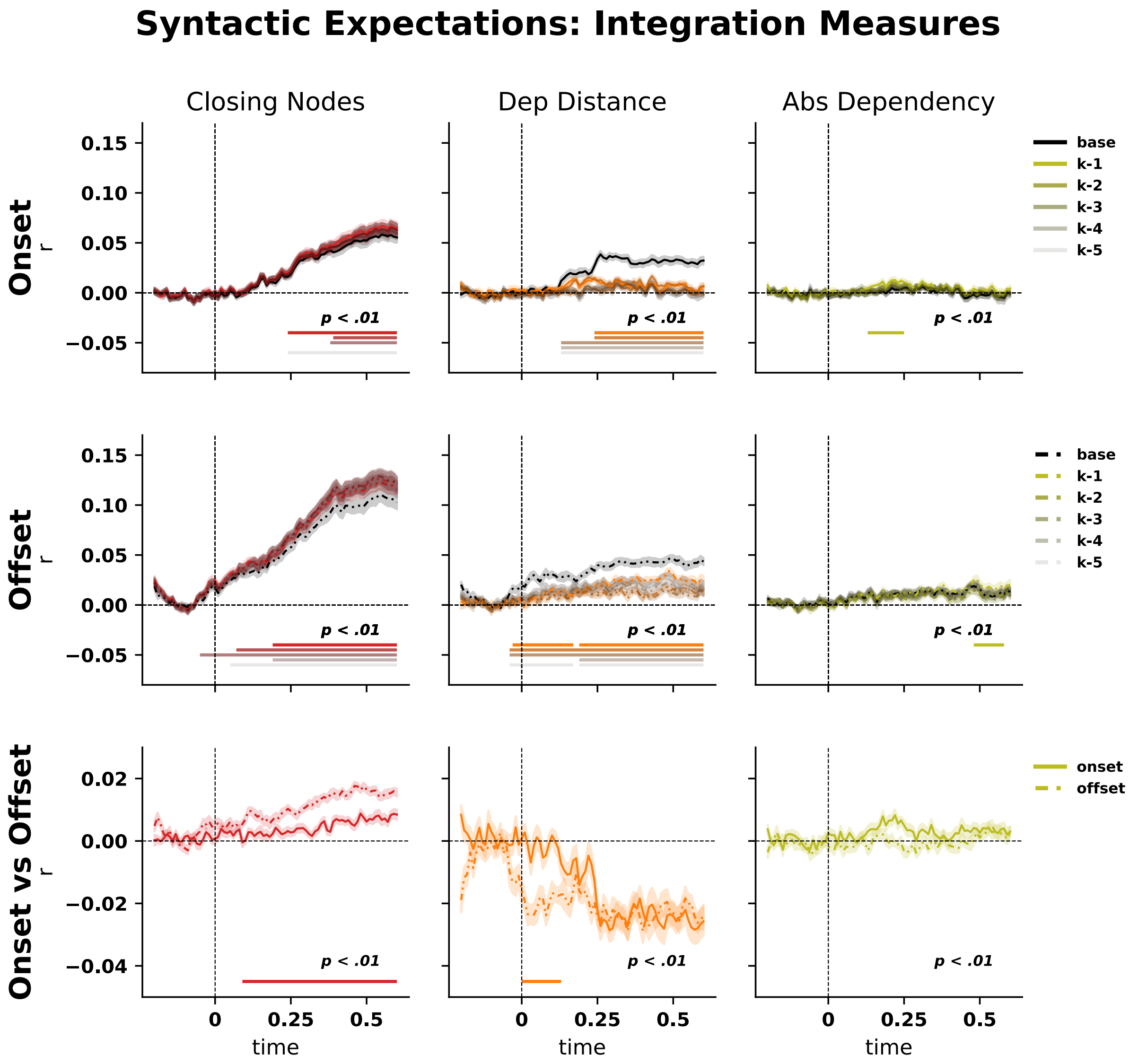


**Supplementary Fig. C 4** Decoding of integration-based syntactic structure using surprisal values. Black lines indicate raw syntactic state feature decoding performance. The significance of color-coded probability measures was computed against baseline vector decoding with a permutation cluster test, and FDR correction ($p_{FDR}$ <.01). Color shading refers to the different contexts, while solid and dashed lines refer to onset and offset respectively. The last row shows the difference between the context -1 and -2 against the base decoding for both onset and offset

while becoming significantly worse at context -4 (0.16 to 0.36 s, $\bar{t}$: -4.1, $p_{\mathrm{FDR}} = \mathit{0.008}$) and context -5 (0.14 to 0.44 s, $\bar{t}$: -4.96, $p_{\mathrm{FDR}} = \mathit{0.008}$).

Integration measures at onset show an enhancement for some measures: closing nodes becomes significantly better at all contexts except context -4 (context -1, 0.24 to 0.6 s, $\bar{t}$: 4.37, $p_{\mathrm{FDR}} = \mathit{0.008}$; context -2, 0.39 to 0.6 s, $\bar{t}$: 3.95, $p_{\mathrm{FDR}} = \mathit{0.008}$; context -3, 0.38 to 0.6 s, $\bar{t}$: 4.19, $p_{\mathrm{FDR}} = \mathit{0.008}$; context -5, 0.24 to 0.6 s, $\bar{t}$: 3.87, $p_{\mathrm{FDR}} = \mathit{0.008}$; no significant change at context -4). *Dependency distance* is significantly worse across contexts (context -1, 0.24 to 0.6 s, $\bar{t}$: -5.78, $p_{\mathrm{FDR}} = \mathit{0.008}$; context -2, 0.24 to 0.6 s, $\bar{t}$: -5.20, $p_{\mathrm{FDR}} = \mathit{0.008}$; context -3, 0.13 to 0.6 s, $\bar{t}$: -6.04, $p_{\mathrm{FDR}} = \mathit{0.008}$; context -4, 0.13 to 0.6 s, $\bar{t}$: -6.01, $p_{\mathrm{FDR}}$

$= 0.008$; context -5, 0.13 to 0.6 s, $\bar{t}$: -6.21, $p_{\mathrm{FDR}} = 0.008$). Lastly, *absolute dependency* becomes significantly better at context -1 (0.13 to 0.25 s, $\bar{t}$: 3.36, $p_{\mathrm{FDR}} = 0.008$) while not changing significantly at all other contexts. Moving to the offset, we observe a similar pattern for memory measures: *tree depth* becomes significantly better at context -1 (-0.06 to 0.6 s, $\bar{t}$: 9.17, $p_{\mathrm{FDR}} = 0.007$), context -2 (-0.06 to 0.6 s, $\bar{t}$: 6.19, $p_{\mathrm{FDR}} = 0.007$), and context -3 (0.12 to 0.18 s, $\bar{t}$: 2.98, $p_{\mathrm{FDR}} = 0.007$). On the contrary, *tree depth* is worse at context -4 (0.35 to 0.6 s, $\bar{t}$: -3.70, $p_{\mathrm{FDR}} = 0.007$) and context -5 (0.34 to 0.6 s, $\bar{t}$: -4.89, $p_{\mathrm{FDR}} = 0.007$). Similarly, *last right* at offset becomes significantly better at context -1 (-0.05 to 0.52 s, $\bar{t}$: 7.54, $p_{\mathrm{FDR}} = 0.007$) and context -2 (0.03 to 0.23 s, $\bar{t}$: 3.92, $p_{\mathrm{FDR}} = 0.007$; 0.26 to 0.42 s, $\bar{t}$: 3.90, $p_{\mathrm{FDR}} = 0.007$), while not changing at context -3. It becomes significantly worse at context -4 (0.18 to 0.6 s, $\bar{t}$: -5.12, $p_{\mathrm{FDR}} = 0.007$) and context -5 (0.18 to 0.6 s, $\bar{t}$: -6.86, $p_{\mathrm{FDR}} = 0.007$). Similarly to the onset counterpart, *dependency depth* at offset shows a enhancement of decoding performance at context -1 (-0.06 to 0.6 s, $\bar{t}$: 5.89, $p_{\mathrm{FDR}} = 0.007$), context -2 (0.26 to 0.6 s, $\bar{t}$: 5.40, $p_{\mathrm{FDR}} = 0.007$), context -3 (0.3 to 0.6 s, $\bar{t}$: 4.12, $p_{\mathrm{FDR}} = 0.007$), and context -4 (0.52 to 0.58 s, $\bar{t}$: 3.59, $p_{\mathrm{FDR}} = 0.007$, while not changing significantly at context -5. For open nodes, context -1 (-0.04 to 0.6 s, $\bar{t}$: 8.98, $p_{\mathrm{FDR}} = 0.007$) and context -2 (-0.04 to 0.6 s, $\bar{t}$: 7.64, $p_{\mathrm{FDR}} = 0.007$) are significantly better, while not changing at other contexts. Finally, integration measures at offset also show a enhancement effect: *closing nodes* is significantly better than base at any context (context -1, 0.19 to 0.6 s, $\bar{t}$: 8.0, $p_{\mathrm{FDR}} = 0.007$; context -2, 0.07 to 0.6 s, $\bar{t}$: 9.86, $p_{\mathrm{FDR}} = 0.007$; context -3, -0.05 to 0.6 s , $\bar{t}$: 9.31, $p_{\mathrm{FDR}} = 0.007$; context -4, 0.19 to 0.6 s, $\bar{t}$: 8.23, $p_{\mathrm{FDR}} = 0.007$; context -5, 0.05 to 0.6 s, $\bar{t}$: 9.75, $p_{\mathrm{FDR}} = 0.007$). As for *dependency distance* at offset, surprisal values show a significant reduction compared to baseline at any context (context -1, -0.03 to 0.17 s, $\bar{t}$: -4.64, $p_{\mathrm{FDR}} = 0.007$; context -1, 0.19 to 0.6 s, $\bar{t}$: -4.82, $p_{\mathrm{FDR}} = 0.007$; context -2, -0.04 to 0.6 s, $\bar{t}$: -6.60, $p_{\mathrm{FDR}} = 0.007$; context -3, -0.04 to 0.6 s, $\bar{t}$: -6.44, $p_{\mathrm{FDR}} = 0.007$; context -4, 0.19 to 0.6 s, $\bar{t}$: -6.53, $p_{\mathrm{FDR}} = 0.007$; context -5, -0.04 to0.17 s, $\bar{t}$: -3.69, $p_{\mathrm{FDR}} = 0.007$; context -5, 0.19 to 0.60 s, $\bar{t}$: -5.26, $p_{\mathrm{FDR}} = 0.007$). Lastly, *absolute dependency* becomes significantly better at context -1 (0.48 to 0.58 s, $\bar{t}$: 3.22, $p_{\mathrm{FDR}} = 0.007$) while not changing significantly at all other contexts. We then compared the sharpening effect between onset and offset by averaging within each measure the results of context -1 and context -2, from which we subtracted the base decoding. We observed an increase for *tree depth* at offset (-0.06 to 0.12 s, $\bar{t}$: -4.53, $p_{\mathrm{FDR}} = 0.006$) and an increase for onset (0.24 to 0.60 s, $\bar{t}$: 3.96, $p_{\mathrm{FDR}} = 0.006$). For last right, we only observed a significant increase for onset (0.23 to 0.60 s $\bar{t}$: 5.97, $p_{\mathrm{FDR}} = 0.006$). *Open nodes* at onset is significantly decreased (0.13 to 0.26 s, $\bar{t}$: -3.76, $p_{\mathrm{FDR}} = 0.006$; 0.32 to 0.60 s, $\bar{t}$: -5.85, $p_{\mathrm{FDR}} = 0.006$). For integration measures, we observed significant changes for *closing nodes* (0.09 to 0.60 s, $\bar{t}$: -4.38, $p_{\mathrm{FDR}} = 0.006$) and *dependency distance* (0.0 to 0.13 s, $\bar{t}$: 3.95, $p_{\mathrm{FDR}} = 0.006$), while not observing any difference for *absolute dependency.*

### Appendix D Generalised sharpening effects

Supplementary Tables D 1, D2, D3 and D4 for memory and integration measures report post hoc contrasts between each probability context and the baseline context, calculated separately for each decoded feature from the full BAM using the model linear-predictor matrix. For each contrast, $p_{\mathrm{raw}}$ is the minimum two-sided pointwise Wald $p$-value across the analysed time course, and $p_{\mathrm{FDR}}$ is the Benjamini-Hochberg-adjusted value obtained across all reported feature-by-context contrasts. Significant time intervals were defined as samples at which the 95% confidence interval of the estimated smooth difference excluded zero.

| contrast | ctx1 | cond1 | ctx2 | cond2 | min_p | p_fdr | significant_fdr_001 |
|---|---|---|---|---|---|---|---|
| prob_1 dependency_depth vs base dependency_depth | prob_1 | dependency_depth | base | dependency_depth | 2.05862994118614e-53 | 3.74296352942935e-53 | TRUE |
| prob_1 last_right vs base last_right | prob_1 | last_right | base | last_right | 4.91152329673859e-98 | 1.63717443224620e-97 | TRUE |
| prob_1 open_nodes vs base open_nodes | prob_1 | open_nodes | base | open_nodes | 1.30304409520757e-26 | 1.86149156458224e-26 | TRUE |
| prob_1 tree_depth vs base tree_depth | prob_1 | tree_depth | base | tree_depth | 1.43423287382046e-124 | 9.56155249213642e-124 | TRUE |
| prob_2 dependency_depth vs base dependency_depth | prob_2 | dependency_depth | base | dependency_depth | 8.54869592661527e-64 | 1.89971020591450e-63 | TRUE |
| prob_2 last_right vs base last_right | prob_2 | last_right | base | last_right | 5.99136022327361e-172 | 1.19827204465472e-170 | TRUE |
| prob_2 open_nodes vs base open_nodes | prob_2 | open_nodes | base | open_nodes | 4.53763647831373e-85 | 1.29646756523249e-84 | TRUE |
| prob_2 tree_depth vs base tree_depth | prob_2 | tree_depth | base | tree_depth | 1.05920150504558e-154 | 1.05920150504558e-153 | TRUE |
| prob_3 dependency_depth vs base dependency_depth | prob_3 | dependency_depth | base | dependency_depth | 2.23235729026919e-21 | 2.97647638702559e-21 | TRUE |
| prob_3 last_right vs base last_right | prob_3 | last_right | base | last_right | 3.13585750692243e-18 | 3.91982188365303e-18 | TRUE |
| prob_3 open_nodes vs base open_nodes | prob_3 | open_nodes | base | open_nodes | 3.08411956920321e-43 | 5.14019928200535e-43 | TRUE |
| prob_3 tree_depth vs base tree_depth | prob_3 | tree_depth | base | tree_depth | 2.72540142555654e-15 | 3.20635461830181e-15 | TRUE |
| prob_4 dependency_depth vs base dependency_depth | prob_4 | dependency_depth | base | dependency_depth | 4.36608678903034e-05 | 4.59588083055825e-05 | TRUE |
| prob_4 last_right vs base last_right | prob_4 | last_right | base | last_right | 3.98284849419518e-59 | 7.96569698839036e-59 | TRUE |
| prob_4 open_nodes vs base open_nodes | prob_4 | open_nodes | base | open_nodes | 5.50289717171332e-11 | 6.11433019079258e-11 | TRUE |
| prob_4 tree_depth vs base tree_depth | prob_4 | tree_depth | base | tree_depth | 3.70107328515211e-71 | 9.25268321288029e-71 | TRUE |
| prob_5 dependency_depth vs base dependency_depth | prob_5 | dependency_depth | base | dependency_depth | 4.69433915233676e-03 | 4.69433915233676e-03 | TRUE |
| prob_5 last_right vs base last_right | prob_5 | last_right | base | last_right | 7.42375524778085e-115 | 3.71187762389043e-114 | TRUE |
| prob_5 open_nodes vs base open_nodes | prob_5 | open_nodes | base | open_nodes | 7.42945514350708e-29 | 1.14299309900109e-28 | TRUE |
| prob_5 tree_depth vs base tree_depth | prob_5 | tree_depth | base | tree_depth | 3.99436851763924e-98 | 1.59774740705570e-97 | TRUE |

**Supplementary Tab. D 1** Post hoc contrasts between probability contexts and baseline for memory-related features at onset.

| contrast | ctx1 | cond1 | ctx2 | cond2 | min_p | p_fdr | significant_fdr_001 |
|---|---|---|---|---|---|---|---|
| prob_1 dependency_depth vs base dependency_depth | prob_1 | dependency_depth | base | dependency_depth | 1.39560150867787e-61 | 3.10133668595083e-61 | TRUE |
| prob_1 last_right vs base last_right | prob_1 | last_right | base | last_right | 1.35504342265454e-102 | 5.42017369061816e-102 | TRUE |
| prob_1 open_nodes vs base open_nodes | prob_1 | open_nodes | base | open_nodes | 0.00000000000000e+00 | 0.00000000000000e+00 | TRUE |
| prob_1 tree_depth vs base tree_depth | prob_1 | tree_depth | base | tree_depth | 1.04607283265419e-179 | 6.97381888436127e-179 | TRUE |
| prob_2 dependency_depth vs base dependency_depth | prob_2 | dependency_depth | base | dependency_depth | 4.12894951847090e-72 | 1.17969986242026e-71 | TRUE |
| prob_2 last_right vs base last_right | prob_2 | last_right | base | last_right | 7.82375915236182e-41 | 1.56475183047236e-40 | TRUE |
| prob_2 open_nodes vs base open_nodes | prob_2 | open_nodes | base | open_nodes | 1.21503538354318e-181 | 1.21503538354318e-180 | TRUE |
| prob_2 tree_depth vs base tree_depth | prob_2 | tree_depth | base | tree_depth | 1.31589837199297e-80 | 4.38632790664325e-80 | TRUE |
| prob_3 dependency_depth vs base dependency_depth | prob_3 | dependency_depth | base | dependency_depth | 1.27302454376867e-34 | 2.31459007957940e-34 | TRUE |
| prob_3 last_right vs base last_right | prob_3 | last_right | base | last_right | 2.88297815712767e-15 | 3.60372269640959e-15 | TRUE |
| prob_3 open_nodes vs base open_nodes | prob_3 | open_nodes | base | open_nodes | 1.43783932742662e-34 | 2.39639887904436e-34 | TRUE |
| prob_3 tree_depth vs base tree_depth | prob_3 | tree_depth | base | tree_depth | 8.05299506552857e-25 | 1.07373267540381e-24 | TRUE |
| prob_4 dependency_depth vs base dependency_depth | prob_4 | dependency_depth | base | dependency_depth | 1.16405012647863e-27 | 1.66292875211233e-27 | TRUE |
| prob_4 last_right vs base last_right | prob_4 | last_right | base | last_right | 3.27098041475019e-32 | 5.03227756115413e-32 | TRUE |
| prob_4 open_nodes vs base open_nodes | prob_4 | open_nodes | base | open_nodes | 2.22435926389387e-02 | 2.22435926389387e-02 | FALSE |
| prob_4 tree_depth vs base tree_depth | prob_4 | tree_depth | base | tree_depth | 4.11497585449578e-15 | 4.84114806411268e-15 | TRUE |
| prob_5 dependency_depth vs base dependency_depth | prob_5 | dependency_depth | base | dependency_depth | 4.96291809427033e-12 | 5.51435343807814e-12 | TRUE |
| prob_5 last_right vs base last_right | prob_5 | last_right | base | last_right | 6.90354106241591e-110 | 3.45177053120796e-109 | TRUE |
| prob_5 open_nodes vs base open_nodes | prob_5 | open_nodes | base | open_nodes | 2.13617532235747e-08 | 2.24860560248155e-08 | TRUE |
| prob_5 tree_depth vs base tree_depth | prob_5 | tree_depth | base | tree_depth | 2.34398645854614e-63 | 5.85996614636536e-63 | TRUE |

**Supplementary Tab. D 2** Post hoc contrasts between probability contexts and baseline for memory-related features at offset.

| contrast | ctx1 | cond1 | ctx2 | cond2 | min_p | p_fdr | significant_fdr_001 |
|---|---|---|---|---|---|---|---|
| prob_1 abs_dependency vs base abs_dependency | prob_1 | abs_dependency | base | abs_dependency | 2.06683709791115e-24 | 2.06683709791115e-24 | TRUE |
| prob_1 closing_nodes vs base closing_nodes | prob_1 | closing_nodes | base | closing_nodes | 7.24010820586831e-300 | 2.17203246176049e-299 | TRUE |
| prob_1 dep_distance vs base dep_distance | prob_1 | dep_distance | base | dep_distance | 4.53809479411058e-73 | 6.80714219116587e-73 | TRUE |
| prob_2 abs_dependency vs base abs_dependency | prob_2 | abs_dependency | base | abs_dependency | 2.22330311942008e-26 | 2.38211048509295e-26 | TRUE |
| prob_2 closing_nodes vs base closing_nodes | prob_2 | closing_nodes | base | closing_nodes | 0.00000000000000e+00 | 0.00000000000000e+00 | TRUE |
| prob_2 dep_distance vs base dep_distance | prob_2 | dep_distance | base | dep_distance | 5.52362630369583e-81 | 9.20604383949305e-81 | TRUE |
| prob_3 abs_dependency vs base abs_dependency | prob_3 | abs_dependency | base | abs_dependency | 2.60257611704510e-42 | 3.25322014630637e-42 | TRUE |
| prob_3 closing_nodes vs base closing_nodes | prob_3 | closing_nodes | base | closing_nodes | 0.00000000000000e+00 | 0.00000000000000e+00 | TRUE |
| prob_3 dep_distance vs base dep_distance | prob_3 | dep_distance | base | dep_distance | 1.67416486913018e-129 | 4.18541217282545e-129 | TRUE |
| prob_4 abs_dependency vs base abs_dependency | prob_4 | abs_dependency | base | abs_dependency | 2.12568030963536e-36 | 2.45270804957926e-36 | TRUE |
| prob_4 closing_nodes vs base closing_nodes | prob_4 | closing_nodes | base | closing_nodes | 0.00000000000000e+00 | 0.00000000000000e+00 | TRUE |
| prob_4 dep_distance vs base dep_distance | prob_4 | dep_distance | base | dep_distance | 1.63486257022502e-112 | 3.50327693619647e-112 | TRUE |
| prob_5 abs_dependency vs base abs_dependency | prob_5 | abs_dependency | base | abs_dependency | 9.63625153548610e-50 | 1.31403430029356e-49 | TRUE |
| prob_5 closing_nodes vs base closing_nodes | prob_5 | closing_nodes | base | closing_nodes | 0.00000000000000e+00 | 0.00000000000000e+00 | TRUE |
| prob_5 dep_distance vs base dep_distance | prob_5 | dep_distance | base | dep_distance | 6.39372121391907e-107 | 1.19882272760983e-106 | TRUE |

**Supplementary Tab. D 3** Post hoc contrasts between probability contexts and baseline for integration-related features at onset.

| contrast | ctx1 | cond1 | ctx2 | cond2 | min_p | p_fdr | significant_fdr_001 |
|---|---|---|---|---|---|---|---|
| prob_1 abs_dependency vs base abs_dependency | prob_1 | abs_dependency | base | abs_dependency | 1.05449734796354e-52 | 1.58174602194530e-51 | TRUE |
| prob_1 closing_nodes vs base closing_nodes | prob_1 | closing_nodes | base | closing_nodes | 2.02132247390864e-21 | 6.06396742172593e-21 | TRUE |
| prob_1 dep_distance vs base dep_distance | prob_1 | dep_distance | base | dep_distance | 3.77191101764728e-21 | 9.42977754411820e-21 | TRUE |
| prob_2 abs_dependency vs base abs_dependency | prob_2 | abs_dependency | base | abs_dependency | 1.45852447882261e-10 | 1.98889701657628e-10 | TRUE |
| prob_2 closing_nodes vs base closing_nodes | prob_2 | closing_nodes | base | closing_nodes | 3.90535379890752e-08 | 4.50617746027790e-08 | TRUE |
| prob_2 dep_distance vs base dep_distance | prob_2 | dep_distance | base | dep_distance | 3.32190437329158e-31 | 2.49142827996869e-30 | TRUE |
| prob_3 abs_dependency vs base abs_dependency | prob_3 | abs_dependency | base | abs_dependency | 3.29951467277150e-05 | 3.53519429225518e-05 | TRUE |
| prob_3 closing_nodes vs base closing_nodes | prob_3 | closing_nodes | base | closing_nodes | 2.90436820658617e-11 | 4.35655230987925e-11 | TRUE |
| prob_3 dep_distance vs base dep_distance | prob_3 | dep_distance | base | dep_distance | 1.09632645634190e-22 | 4.11122421128213e-22 | TRUE |
| prob_4 abs_dependency vs base abs_dependency | prob_4 | abs_dependency | base | abs_dependency | 3.74890597551556e-16 | 8.03336994753334e-16 | TRUE |
| prob_4 closing_nodes vs base closing_nodes | prob_4 | closing_nodes | base | closing_nodes | 1.29521307792900e-12 | 2.15868846321501e-12 | TRUE |
| prob_4 dep_distance vs base dep_distance | prob_4 | dep_distance | base | dep_distance | 1.16414891717429e-09 | 1.45518614646786e-09 | TRUE |
| prob_5 abs_dependency vs base abs_dependency | prob_5 | abs_dependency | base | abs_dependency | 4.10781426495043e-26 | 2.05390713247522e-25 | TRUE |
| prob_5 closing_nodes vs base closing_nodes | prob_5 | closing_nodes | base | closing_nodes | 3.44035489001777e-14 | 6.45066541878331e-14 | TRUE |
| prob_5 dep_distance vs base dep_distance | prob_5 | dep_distance | base | dep_distance | 1.85927835061799e-04 | 1.85927835061799e-04 | TRUE |

**Supplementary Tab. D 4** Post hoc contrasts between probability contexts and baseline for integration-related features at offset.

### Appendix E Inverted U-shape

We include separate quadratic-model tables (Supplementary Tab. E 1 and E 2) for probability and surprisal measures. These analyses tested whether decoding across probability contexts followed a significant quadratic profile, separately for each decoded feature and temporal phase. For each feature-by-phase combination, the table reports the linear and quadratic coefficients, the raw and FDR-corrected p-values for the quadratic term, the estimated turning point, and whether that turning point fell within the observed context range. The resulting classification indicates whether the profile was a significant U-shape, a significant inverted-U-shape, a significant quadratic effect with the turning point outside the sampled range, or no significant quadratic shape.

| condition | phase | beta_linear | p_linear | beta_quadratic | p_quadratic | p_quadratic_fdr | turning_point_centered | turning_point_original | turning_point_in_range | shape |
|---|---|---|---|---|---|---|---|---|---|---|
| abs_dependency | offset | 6.43639e-04 | 5.16888e-03 | -9.25047e-05 | 5.51304e-01 | 5.51304e-01 | 3.4789538 | 6.97895 | FALSE | no significant quadratic shape |
| closing_nodes | offset | -1.02668e-03 | 3.78112e-10 | 5.78950e-04 | 1.26475e-07 | 1.47555e-07 | 0.8866723 | 4.38667 | TRUE | significant U-shape |
| dep_distance | offset | -3.93440e-04 | 4.44219e-02 | 1.99086e-03 | 1.06104e-29 | 7.42729e-29 | 0.0988114 | 3.59881 | TRUE | significant U-shape |
| dependency_depth | offset | -2.30989e-04 | 4.24540e-01 | -1.78862e-03 | 1.18283e-15 | 1.83996e-15 | -0.0645720 | 3.43543 | TRUE | significant inverted U-shape |
| last_right | offset | -5.28797e-03 | 2.13722e-26 | -3.63959e-03 | 1.43132e-26 | 6.67951e-26 | -0.7264515 | 2.77355 | TRUE | significant inverted U-shape |
| open_nodes | offset | -4.53923e-03 | 1.05725e-10 | -4.37570e-03 | 9.39939e-18 | 1.64489e-17 | -0.5186864 | 2.98131 | TRUE | significant inverted U-shape |
| tree_depth | offset | -5.48866e-03 | 1.34106e-19 | -4.14404e-03 | 2.24098e-22 | 5.22894e-22 | -0.6622357 | 2.83776 | TRUE | significant inverted U-shape |
| abs_dependency | onset | -6.27353e-04 | 3.19280e-06 | -1.43030e-04 | 1.06986e-01 | 1.15216e-01 | -2.1930818 | 1.30692 | TRUE | no significant quadratic shape |
| closing_nodes | onset | -1.76086e-03 | 2.85316e-25 | 7.93804e-04 | 2.08150e-14 | 2.91410e-14 | 1.1091310 | 4.60913 | TRUE | significant U-shape |
| dep_distance | onset | -2.93151e-03 | 5.82740e-30 | 1.07134e-03 | 6.52280e-13 | 8.30174e-13 | 1.3681469 | 4.86815 | TRUE | significant U-shape |
| dependency_depth | onset | -1.46370e-03 | 6.01059e-06 | -2.76913e-03 | 1.65776e-25 | 5.80217e-25 | -0.2642881 | 3.23571 | TRUE | significant inverted U-shape |
| last_right | onset | -7.85135e-03 | 1.27681e-19 | -6.51668e-03 | 2.56979e-25 | 7.19540e-25 | -0.6024045 | 2.89760 | TRUE | significant inverted U-shape |
| open_nodes | onset | -2.81078e-03 | 2.20136e-09 | -4.82510e-03 | 6.62097e-33 | 9.26936e-32 | -0.2912658 | 3.20873 | TRUE | significant inverted U-shape |
| tree_depth | onset | -7.83984e-03 | 1.19215e-19 | -5.78096e-03 | 9.85752e-22 | 1.97150e-21 | -0.6780748 | 2.82193 | TRUE | significant inverted U-shape |

**Supplementary Tab. E 1** Quadratic context-profile tests for probability measures

| condition | phase | beta_linear | p_linear | beta_quadratic | p_quadratic | p_quadratic_fdr | turning_point_centered | turning_point_original | turning_point_in_range | shape |
|---|---|---|---|---|---|---|---|---|---|---|
| abs_dependency | offset | 6.43639e-04 | 5.16888e-03 | -9.25047e-05 | 5.51304e-01 | 5.51304e-01 | 3.4789538 | 6.97895 | FALSE | no significant quadratic shape |
| closing_nodes | offset | -1.02668e-03 | 3.78112e-10 | 5.78950e-04 | 1.26475e-07 | 1.47555e-07 | 0.8866723 | 4.38667 | TRUE | significant U-shape |
| dep_distance | offset | -3.93440e-04 | 4.44219e-02 | 1.99086e-03 | 1.06104e-29 | 7.42729e-29 | 0.0988114 | 3.59881 | TRUE | significant U-shape |
| dependency_depth | offset | -2.30989e-04 | 4.24540e-01 | -1.78862e-03 | 1.18283e-15 | 1.83996e-15 | -0.0645720 | 3.43543 | TRUE | significant inverted U-shape |
| last_right | offset | -5.28797e-03 | 2.13722e-26 | -3.63959e-03 | 1.43132e-26 | 6.67951e-26 | -0.7264515 | 2.77355 | TRUE | significant inverted U-shape |
| open_nodes | offset | -4.53923e-03 | 1.05725e-10 | -4.37570e-03 | 9.39939e-18 | 1.64489e-17 | -0.5186864 | 2.98131 | TRUE | significant inverted U-shape |
| tree_depth | offset | -5.48866e-03 | 1.34106e-19 | -4.14404e-03 | 2.24098e-22 | 5.22894e-22 | -0.6622357 | 2.83776 | TRUE | significant inverted U-shape |
| abs_dependency | onset | -6.27353e-04 | 3.19280e-06 | -1.43030e-04 | 1.06986e-01 | 1.15216e-01 | -2.1930818 | 1.30692 | TRUE | no significant quadratic shape |
| closing_nodes | onset | -1.76086e-03 | 2.85316e-25 | 7.93804e-04 | 2.08150e-14 | 2.91410e-14 | 1.1091310 | 4.60913 | TRUE | significant U-shape |
| dep_distance | onset | -2.93151e-03 | 5.82740e-30 | 1.07134e-03 | 6.52280e-13 | 8.30174e-13 | 1.3681469 | 4.86815 | TRUE | significant U-shape |
| dependency_depth | onset | -1.46370e-03 | 6.01059e-06 | -2.76913e-03 | 1.65776e-25 | 5.80217e-25 | -0.2642881 | 3.23571 | TRUE | significant inverted U-shape |
| last_right | onset | -7.85135e-03 | 1.27681e-19 | -6.51668e-03 | 2.56979e-25 | 7.19540e-25 | -0.6024045 | 2.89760 | TRUE | significant inverted U-shape |
| open_nodes | onset | -2.81078e-03 | 2.20136e-09 | -4.82510e-03 | 6.62097e-33 | 9.26936e-32 | -0.2912658 | 3.20873 | TRUE | significant inverted U-shape |
| tree_depth | onset | -7.83984e-03 | 1.19215e-19 | -5.78096e-03 | 9.85752e-22 | 1.97150e-21 | -0.6780748 | 2.82193 | TRUE | significant inverted U-shape |

**Supplementary Tab. E 2** Quadratic context-profile tests for surprisal measures

### Appendix F Anchoring probability decoding

We include the model-comparison figures and tables Supplementary Fig. F 1 and F 2, Supplementary Tab. F 1 and F 2, for Memory measures and Integration measures respectively, which report on the temporal anchoring effect (alignment to word onset or offset), based on model comparisons within each condition-by- probability context cell, and comparing the reduced BAM with the full BAM using a chi-square test. Resulting figures were plotted using `ggplot2` [2]. Here, each *p_raw* denotes the p value for the overall model comparison.

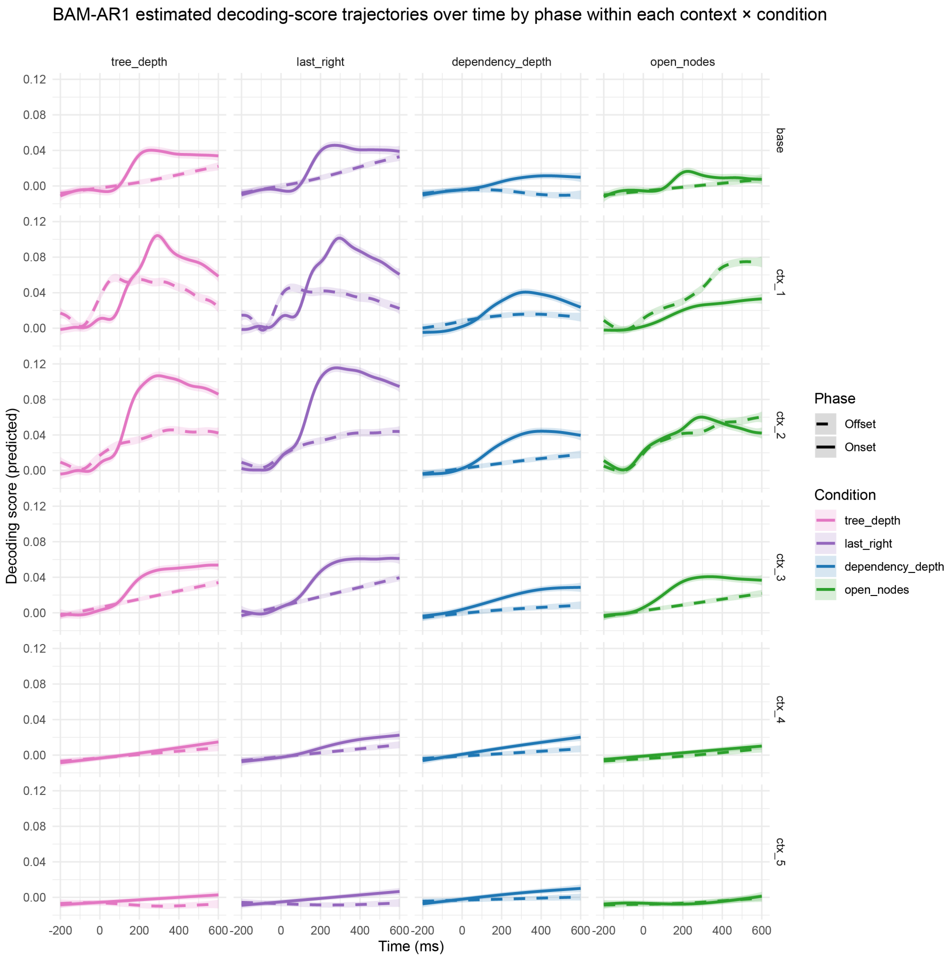


**Supplementary Fig. F 1** BAM-AR1 estimated decoding-score trajectories for memory-related syntactic features across contexts. Predicted decoding trajectories are shown separately for onset- and offset-aligned analyses in each context × condition cell. Solid lines indicate onset and dashed lines offset; shaded ribbons represent 95% confidence intervals. Colors denote syntactic features.

| condition | context | rho | Df | Deviance | p_raw | p_fdr | significant_fdr_01 |
|---|---|---|---|---|---|---|---|
| dependency_depth | base | 0.50380499059562 | 13.9536009568974 | 0.101271777069374 | 2.01388742188888e-113 | 3.40811717550412e-113 | TRUE |
| dependency_depth | ctx_1 | 0.600419496043578 | 22.3305491055335 | 0.152639402173471 | 1.8119617186452e-130 | 3.62392343729039e-130 | TRUE |
| dependency_depth | ctx_2 | 0.579256139210252 | 1.14250215105812 | 0.160206370304695 | 8.20316342530135e-171 | 2.00521772618477e-170 | TRUE |
| dependency_depth | ctx_3 | 0.518627636811521 | 2.42848663542009 | 0.0586069806614495 | 2.53985050161868e-67 | 3.99119364540078e-67 | TRUE |
| dependency_depth | ctx_4 | 0.448513297738917 | 0.165540065566802 | 0.0208089632458829 | 1.00859717183194e-29 | 1.30524339884133e-29 | TRUE |
| dependency_depth | ctx_5 | 0.441948284086855 | 14.623232079136 | 0.0184969459386212 | 2.08989287766989e-16 | 2.29888216543688e-16 | TRUE |
| last_right | base | 0.61122378908046 | 8.82254249554489 | 0.196133904359492 | 3.05833751694597e-186 | 8.41042817160142e-186 | TRUE |
| last_right | ctx_1 | 0.830943013511995 | 1.41174590750006 | 0.995888445853386 | 9.99999999999999e-301 | 9.99999999999999e-301 | TRUE |
| last_right | ctx_2 | 0.837336953764272 | 4.62730908250296 | 1.10035408176453 | 9.99999999999999e-301 | 9.99999999999999e-301 | TRUE |
| last_right | ctx_3 | 0.619720979460725 | 7.08934689549505 | 0.210126862217201 | 6.03924628486822e-193 | 1.89804883238715e-192 | TRUE |
| last_right | ctx_4 | 0.461282667781772 | 4.97559828401108 | 0.0295119269746474 | 2.87228907030252e-34 | 4.2126906364437e-34 | TRUE |
| last_right | ctx_5 | 0.443599193474668 | 3.40265328217674 | 0.0259294922158384 | 8.38754107392274e-32 | 1.15328689766438e-31 | TRUE |
| open_nodes | base | 0.473239894680074 | 8.32234666687009 | 0.0217147349360248 | 1.23480672617844e-21 | 1.42977620925924e-21 | TRUE |
| open_nodes | ctx_1 | 0.66201240634514 | 5.34075318469922 | 0.24885383445257 | 6.20567955750811e-217 | 2.73049900530357e-216 | TRUE |
| open_nodes | ctx_2 | 0.622047411256519 | -0.635293290162281 | 0.0620521275848 | NA | NA | NA |
| open_nodes | ctx_3 | 0.550385880563548 | 4.07084615341682 | 0.109397600222609 | 2.48005881453302e-114 | 4.54677449331053e-114 | TRUE |
| open_nodes | ctx_4 | 0.440994621933489 | 4.31614232809488 | 0.00198910083634407 | 0.0275822174359168 | 0.0275822174359168 | FALSE |
| open_nodes | ctx_5 | 0.408409297139196 | 4.85155360782619 | -0.000479427863383508 | NA | NA | NA |
| tree_depth | base | 0.600703818225679 | 10.1473247861641 | 0.203286429906775 | 2.14308859997525e-196 | 7.85799153324259e-196 | TRUE |
| tree_depth | ctx_1 | 0.814511307004448 | 2.32865143965546 | 0.880544196826105 | 9.99999999999999e-301 | 9.99999999999999e-301 | TRUE |
| tree_depth | ctx_2 | 0.812893673334651 | 6.26807970416257 | 0.91913081397226 | 9.99999999999999e-301 | 9.99999999999999e-301 | TRUE |
| tree_depth | ctx_3 | 0.598504228973619 | 6.39092678783982 | 0.167352389663184 | 1.82835582608839e-165 | 4.02238281739447e-165 | TRUE |
| tree_depth | ctx_4 | 0.433093685766666 | 8.29317908841494 | 0.00912894232271311 | 7.81094524519339e-09 | 8.18289501877403e-09 | TRUE |
| tree_depth | ctx_5 | 0.425459343129891 | 4.20780903728291 | 0.0193673205251965 | 2.50539682798702e-23 | 3.0621516786508e-23 | TRUE |

**Supplementary Tab. F 1** Supplementary Table for temporal anchor effect model comparisons for Memory features using probability contexts. NA indicates that the reduced-versus-full GAMM comparison yielded a negative effective degrees-of-freedom difference, so an interpretable chi-square p value was not available.

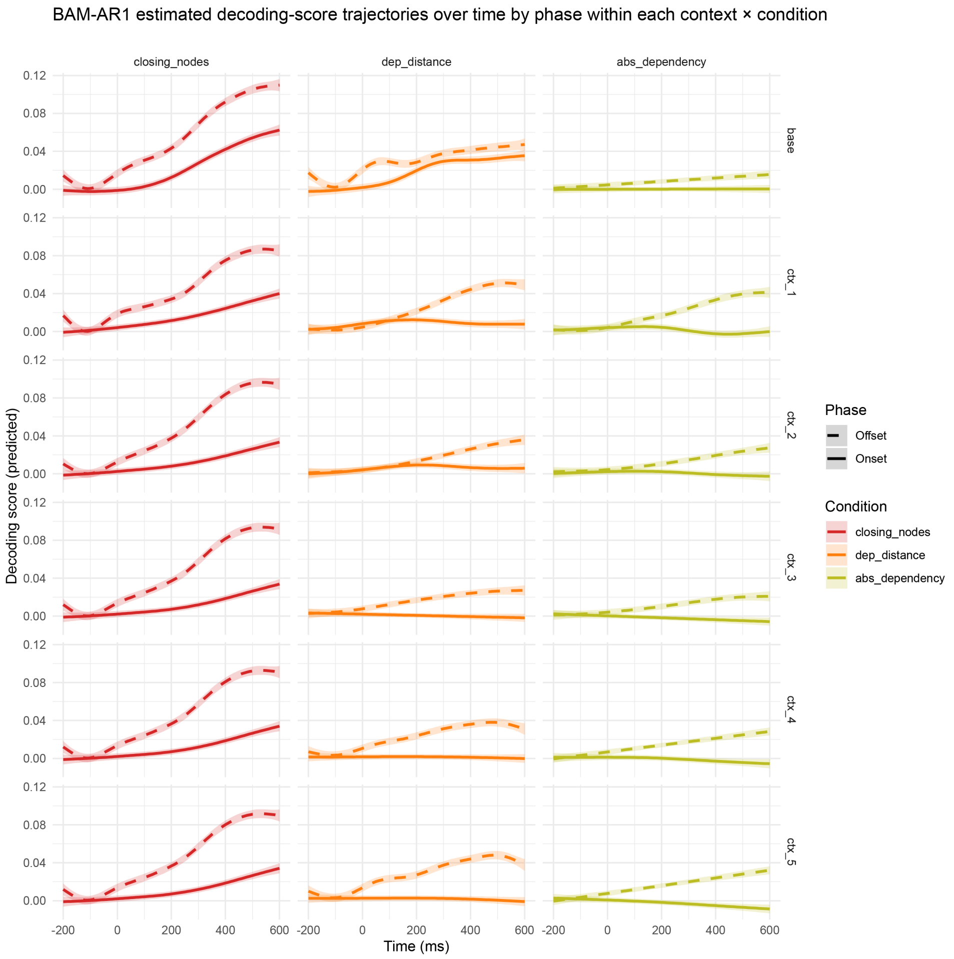


**Supplementary Fig. F 2** BAM-AR1 estimated decoding-score trajectories for integration-related syntactic features across contexts. Predicted decoding trajectories are shown separately for onset- and offset-aligned analyses in each context × condition cell. Solid lines indicate onset and dashed lines offset; shaded ribbons represent 95% confidence intervals. Colors denote syntactic features.

| condition | context | rho | Df | Deviance | p_raw | p_fdr | significant_fdr_01 |
|---|---|---|---|---|---|---|---|
| abs_dependency | base | 0.509556245972665 | 5.39866570393451 | 0.0167606694367299 | 1.8768124896564e-17 | 1.8768124896564e-17 | TRUE |
| abs_dependency | ctx_1 | 0.643127955719655 | 9.21806725137003 | 0.298298661750321 | 3.21465847739972e-251 | 8.26626465617071e-251 | TRUE |
| abs_dependency | ctx_2 | 0.562994760348066 | 19.8465188714413 | 0.106142446766498 | 5.50316058898727e-93 | 6.19105566261068e-93 | TRUE |
| abs_dependency | ctx_3 | 0.541914662894583 | 7.89093589777849 | 0.0992043570439887 | 7.95864208214297e-98 | 9.55037049857156e-98 | TRUE |
| abs_dependency | ctx_4 | 0.566136036108022 | 6.45046201992773 | 0.128672753740147 | 3.04539774813293e-119 | 4.5680966221994e-119 | TRUE |
| abs_dependency | ctx_5 | 0.593494052449058 | 3.30270602426936 | 0.184366711553726 | 2.17664942597014e-171 | 3.91796896674626e-171 | TRUE |
| closing_nodes | base | 0.698805844258276 | 7.72120530490974 | 0.277193056998721 | 2.3162035557759e-208 | 4.63240711155181e-208 | TRUE |
| closing_nodes | ctx_1 | 0.682051007017236 | 11.9090500659145 | 0.33779405112281 | 5.03381456979439e-263 | 1.81217324512598e-262 | TRUE |
| closing_nodes | ctx_2 | 0.754255246721052 | 3.88235525394748 | 0.640203954821857 | 9.99999999999999e-301 | 9.99999999999999e-301 | TRUE |
| closing_nodes | ctx_3 | 0.73885228373914 | 5.90012924956136 | 0.577071635114269 | 9.99999999999999e-301 | 9.99999999999999e-301 | TRUE |
| closing_nodes | ctx_4 | 0.734409040270486 | 5.67476606976061 | 0.552909115434395 | 9.99999999999999e-301 | 9.99999999999999e-301 | TRUE |
| closing_nodes | ctx_5 | 0.730588519126278 | 6.76469397313167 | 0.537000291533137 | 9.99999999999999e-301 | 9.99999999999999e-301 | TRUE |
| dep_distance | base | 0.584308821003454 | 7.90808998225157 | 0.0377159311002873 | 5.57438899265014e-34 | 5.90229422751191e-34 | TRUE |
| dep_distance | ctx_1 | 0.67897640605925 | 5.35381106357318 | 0.324651626529503 | 8.88862000586268e-260 | 2.6665860017588e-259 | TRUE |
| dep_distance | ctx_2 | 0.580250063210989 | 9.40541010155903 | 0.124766577737295 | 1.09188064553155e-115 | 1.51183473996677e-115 | TRUE |
| dep_distance | ctx_3 | 0.543686932753776 | 0.13002463358589 | 0.105663307339924 | 4.24022989703908e-115 | 5.45172415333596e-115 | TRUE |
| dep_distance | ctx_4 | 0.589843780035486 | 5.70372205454805 | 0.157575987303311 | 2.07636382351932e-146 | 3.39768625666798e-146 | TRUE |
| dep_distance | ctx_5 | 0.650463511847206 | 7.72790847658553 | 0.248540615203852 | 8.86055062851874e-209 | 1.99362389141672e-208 | TRUE |

**Supplementary Tab. F 2** Supplementary Table for temporal anchor effect model comparisons for Integration features using probability contexts

### Appendix G Anchoring surprisal decoding

We tested the trajectories of surprisal decoding profiles by fitting again an omnibus subject × condition (decoding feature) × context (probability bin) × word anchor (onset, offset) General Additive Mixed Model (GAMM), separately for memory and integration feature groups. The omnibus model included an AR1 error structure to control for the inflating effects of residual autocorrelation across adjacent time points (see Methods). The full model fit the data better than the reduced model, highlighing significant word anchor × context × condition combination : memory measures, $\Delta\chi^2 = 5.656$, $\Delta\text{df} = 129.21$, $p =$ 2.2e-16 ; integration measures, $\Delta\chi^2 = 2.602$, $\Delta\text{df} = 65.772$, $p =$ 2.2e-16. Thus, also for suprisal selecting a temporal anchor for alignment changes decoding performance. For memory measures, positive onset–offset contrasts indicated generally stronger decoding at onset than at offset across all memory measures, with all context × condition contrasts surviving FDR correction. The largest effects were observed for *last right*, particularly at lags $-1$ and $-2$, followed by *tree depth*, which showed a strong onset advantage across both nearby and more distant contexts. *Dependency depth* displayed a smaller but reliable positive effect that declined with contextual distance, whereas *open nodes* showed a weaker and less locally concentrated pattern (see Supplementary Tab. G 1). For integration measures, negative onset–offset contrasts indicated generally stronger decoding at offset than at onset across all integration measures, with all context × condition contrasts surviving FDR correction. The strongest effects were observed for *closing nodes*, whose offset advantage was large and consistent across all contexts. *Dependency distance* showed a more moderate but still reliable negative effect, with a less uniform profile across contextual distance. *Absolute dependency* exhibited the smallest offset advantage overall, although this effect remained significant throughout the full context range (see Supplementary Tab. G 2). In all cases, onset–offset differences extended across much of the trajectory, confirming also for surprisal that alignment effects reflect a stable property of decoding dynamics rather than residual carry-over between adjacent samples. This pattern further supports the theoretical distinction between memory and integration measures.

We include below the model-comparison tables, for Memory measures and Integration measures respectively, which report on the temporal anchoring effect (alignment to word onset or offset), based on model comparisons within each condition-by- surprisal context cell, and comparing the reduced BAM with the full BAM using a chi-square test. Each *p_raw* denotes the p value for the overall model comparison.

### Appendix H Low-level confounds

To address potential confounds, we report the relation between low-level features and our syntactic features. For low-level features we included word frequency, derived from the Python package `wordfreq` [1], word length computed as the number of characters in the each word, and word index, provided

| condition | context | rho | Df | Deviance | p_raw | p_fdr | significant_fdr_01 |
|---|---|---|---|---|---|---|---|
| dependency_depth | base | 0.50380499059562 | 13.9536009568974 | 0.101271777069374 | 2.01388742188888e-113 | 3.84469416906043e-113 | TRUE |
| dependency_depth | ctx_1 | 0.686485517779096 | 5.22521259408722 | 0.311569241049101 | 3.32761406859161e-233 | 1.39759790880848e-232 | TRUE |
| dependency_depth | ctx_2 | 0.52568781671899 | 6.6698635809571 | 0.0772312174432631 | 1.13222016495186e-86 | 1.98138528866575e-86 | TRUE |
| dependency_depth | ctx_3 | 0.479218414646123 | 5.30926398658175 | 0.00787857933999236 | 3.07530224264047e-08 | 3.07530224264047e-08 | TRUE |
| dependency_depth | ctx_4 | 0.444342952691584 | 3.5471780660182 | 0.00841248320301913 | 2.17028023572519e-10 | 2.39873078685416e-10 | TRUE |
| dependency_depth | ctx_5 | 0.436484303952306 | 1.89390644820651 | 0.0200427306952278 | 1.35647043156542e-26 | 2.03470564734813e-26 | TRUE |
| last_right | base | 0.61122378908046 | 8.82378444657752 | 0.195976910097423 | 4.44212007408842e-186 | 1.16605651944821e-185 | TRUE |
| last_right | ctx_1 | 0.863477734250275 | 8.89610499480932 | 1.32479375928119 | 9.99999999999999e-301 | 9.99999999999999e-301 | TRUE |
| last_right | ctx_2 | 0.770942660788676 | 15.7907112429298 | 0.630427997865282 | 9.99999999999999e-301 | 9.99999999999999e-301 | TRUE |
| last_right | ctx_3 | 0.559813493028244 | 4.23046801629744 | 0.128852339560981 | 2.45834519733898e-139 | 5.16252491441186e-139 | TRUE |
| last_right | ctx_4 | 0.441366810908358 | 4.12220457548437 | 0.0188951776871031 | 3.08799803750162e-22 | 4.05299742422087e-22 | TRUE |
| last_right | ctx_5 | 0.414481028363202 | 4.51473386962562 | 0.0196944538681091 | 9.75743927091115e-24 | 1.36604149792756e-23 | TRUE |
| open_nodes | base | 0.473239894680074 | 8.32234666686963 | 0.0217147349360242 | 1.23480672617977e-21 | 1.52534948528089e-21 | TRUE |
| open_nodes | ctx_1 | 0.647935898558871 | 9.0701807695259 | 0.231278729376025 | 1.77538338748131e-200 | 6.21384185618457e-200 | TRUE |
| open_nodes | ctx_2 | 0.537672497239622 | 10.5549904061318 | 0.0370452391220273 | 9.21005560819224e-36 | 1.48777821363105e-35 | TRUE |
| open_nodes | ctx_3 | 0.46103017479572 | -7.53113001929842 | 0.0439077303919172 | NA | NA | NA |
| open_nodes | ctx_4 | 0.391419141639084 | 4.23629656603725 | 0.0073375504797295 | 6.87961401529541e-09 | 7.22359471606018e-09 | TRUE |
| open_nodes | ctx_5 | 0.390047175533083 | 2.25697696530096 | -0.00185179875322905 | NA | NA | NA |
| tree_depth | base | 0.600703818225679 | 10.1473247861641 | 0.203286429906775 | 2.14308859997525e-196 | 6.42926579992575e-196 | TRUE |
| tree_depth | ctx_1 | 0.86216080141774 | 9.45403287248291 | 1.52038023560294 | 9.99999999999999e-301 | 9.99999999999999e-301 | TRUE |
| tree_depth | ctx_2 | 0.753333395955354 | 11.6563231838331 | 0.587576474926811 | 9.99999999999999e-301 | 9.99999999999999e-301 | TRUE |
| tree_depth | ctx_3 | 0.564715488536017 | 3.94858834495153 | 0.135697918077641 | 5.23806357505388e-142 | 1.22221483417924e-141 | TRUE |
| tree_depth | ctx_4 | 0.429545599003681 | -1.07306972407287 | 0.0159031753060592 | NA | NA | NA |
| tree_depth | ctx_5 | 0.448906013982806 | 0.292222698867135 | 0.0088355938354755 | 8.41482423027426e-14 | 9.81729493531997e-14 | TRUE |

**Supplementary Tab. G 1** Supplementary Table for temporal anchor effect model comparisons for Memory features using surprisal contexts. NA indicates that the reduced-versus-full GAMM comparison yielded a negative effective degrees-of-freedom difference, so an interpretable chi-square p value was not available.

| condition | context | rho | Df | Deviance | p_raw | p_fdr | significant_fdr_01 |
|---|---|---|---|---|---|---|---|
| abs_dependency | base | 0.509557301846549 | -2.8260992794294 | 0.0154487512220312 | NA | NA | NA |
| abs_dependency | ctx_1 | 0.520503897489563 | 6.31707305004602 | 0.0339811647011757 | 1.031041993137e-34 | 2.062083986274e-34 | TRUE |
| abs_dependency | ctx_2 | 0.47548056924691 | 5.35365013388218 | 0.00921250452676503 | 1.9038963877997e-09 | 2.34325709267656e-09 | TRUE |
| abs_dependency | ctx_3 | 0.464742135890434 | 2.00908762179552 | 0.00397146066531029 | 1.57844849567286e-05 | 1.68367839538438e-05 | TRUE |
| abs_dependency | ctx_4 | 0.482379295709773 | 5.39886254785461 | 0.0167086202112774 | 9.88927211677236e-18 | 1.31856961556965e-17 | TRUE |
| abs_dependency | ctx_5 | 0.493765732448445 | 20.3752463755136 | 0.0244739622700781 | 2.76823915230618e-18 | 4.02652967608172e-18 | TRUE |
| closing_nodes | base | 0.698783547904314 | 7.13267613056678 | 0.271326125486395 | 2.15141270385363e-204 | 5.73710054360969e-204 | TRUE |
| closing_nodes | ctx_1 | 0.724529904643728 | 7.96215522359944 | 0.323350678164656 | 2.54269456021326e-221 | 8.13662259268245e-221 | TRUE |
| closing_nodes | ctx_2 | 0.752984802318417 | 7.94087149348752 | 0.456244312756526 | 4.76439704205533e-285 | 3.81151763364427e-284 | TRUE |
| closing_nodes | ctx_3 | 0.749842277246389 | 8.67268840770521 | 0.45689501561522 | 4.25592406756923e-285 | 3.81151763364427e-284 | TRUE |
| closing_nodes | ctx_4 | 0.743801139922153 | 18.8568794694452 | 0.432603313671388 | 2.92187210469875e-263 | 1.1687488418795e-262 | TRUE |
| closing_nodes | ctx_5 | 0.74936291270844 | 8.59230213965611 | 0.425210247687961 | 4.78616380207384e-266 | 2.55262069443938e-265 | TRUE |
| dep_distance | base | 0.584380837498446 | 4.65331377438406 | 0.0333107494585704 | 4.52810408056778e-32 | 8.04996280989828e-32 | TRUE |
| dep_distance | ctx_1 | 0.543794191799108 | -0.906896273541861 | 0.0605747692529633 | NA | NA | NA |
| dep_distance | ctx_2 | 0.454029052265094 | 2.85259346872317 | 0.00569907787963253 | 3.7317155655504e-07 | 4.26481778920046e-07 | TRUE |
| dep_distance | ctx_3 | 0.457832313258601 | 4.31475524826192 | 0.00466020559802838 | 3.7751691306468e-05 | 3.7751691306468e-05 | TRUE |
| dep_distance | ctx_4 | 0.474863620869473 | 7.96673191061291 | 0.0187000532019478 | 8.73329538984107e-19 | 1.39732726237457e-18 | TRUE |
| dep_distance | ctx_5 | 0.524953504244339 | 19.9007166444867 | 0.0459094047081147 | 1.46929828096361e-39 | 3.35839607077398e-39 | TRUE |

**Supplementary Tab. G 2** Supplementary Table for temporal anchor effect model comparisons for Integration features using surprisal contexts. NA indicates that the reduced-versus-full GAMM comparison yielded a negative effective degrees-of-freedom difference, so an interpretable chi-square p value was not available.

as part of the annotations in the dataset. Here we report the full correlation matrix between these low-level confounds and syntactic features (Supplementary Fig. H 1 and Supplementary Fig. H 2). In Supplementary Fig. H2 we also report the full correlation matrix of syntactic features and their marginalized transition probabilities.

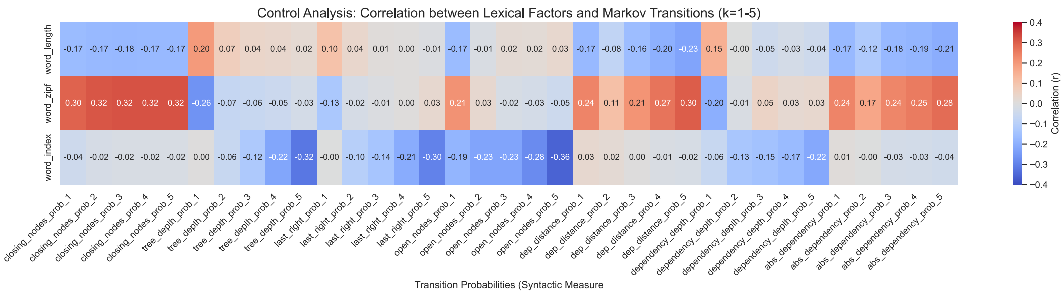


**Supplementary Fig. H 1** Correlation matrix between probability variables and low-level features (word index in the sentence, word length, and word frequency. Word Frequency was added from the wordfreq package in Python.

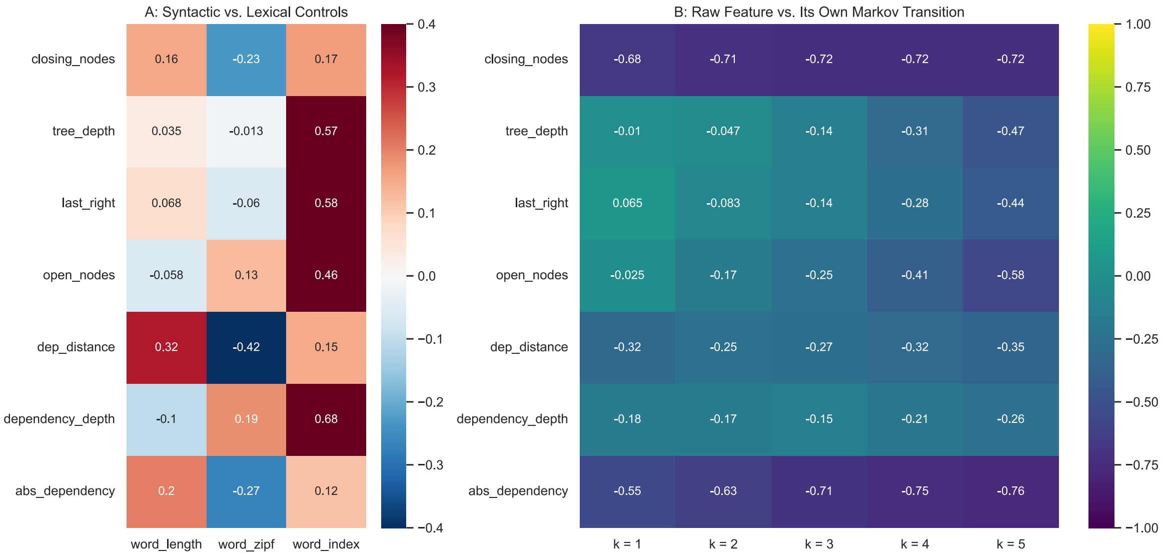


**Supplementary Fig. H 2** Figure A shows the correlation matrix between raw syntactic features and low-level features. As expected, we observe a high correlation for word index, but also with word length for some of the features; Figure B shows the correlation matrix of each syntactic feature and its own marginalized transition probabilities

### Appendix I Partial Correlations

We evaluated the impact of low-level features by running a partial correlation analysis. As in our main analysis, we fit the decoding model to the MEG data and generate predicted values in cross validation (K = 5), separately for session 1 and session 2. Before correlating the predicted values with the ground-truth values, we partial out word frequency, word length, and word index. Here we report the figure and the permutation cluster tests for Memory and Integration measures (see Supplementary Fig. I 1 and Supplementary Fig. I 2). We observe that the overall pattern of decoding performance is maintained across both group of measures.

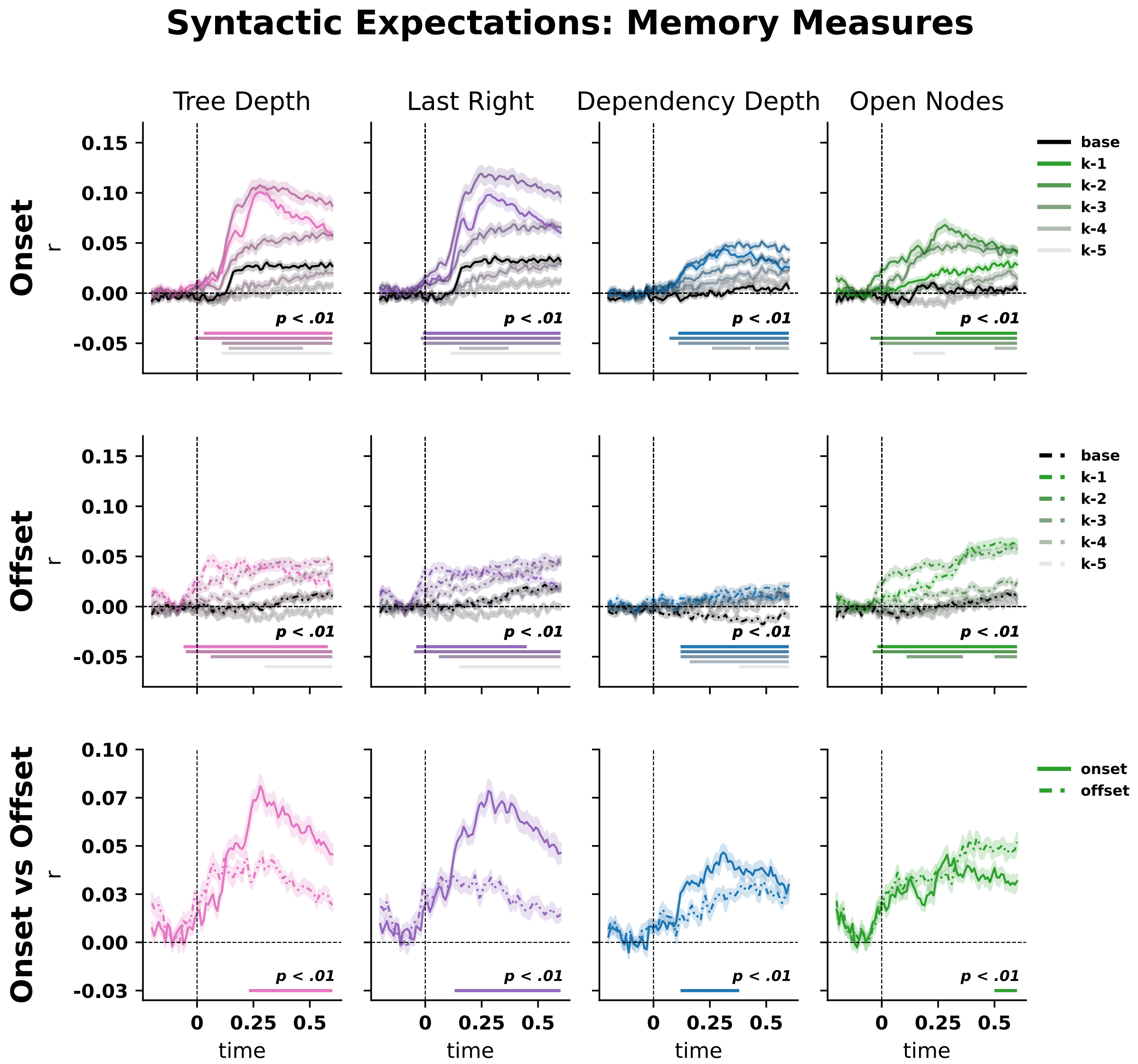


**Supplementary Fig. I 1** Partial correlation analysis of memory features. Before evaluating the decoding model's prediction we residualise our true variable and predicted variable and then compute the Pearson correlation. For memory measure we observe a reduction of the raw features, which allows context -3 to be also significantly better than baseline.

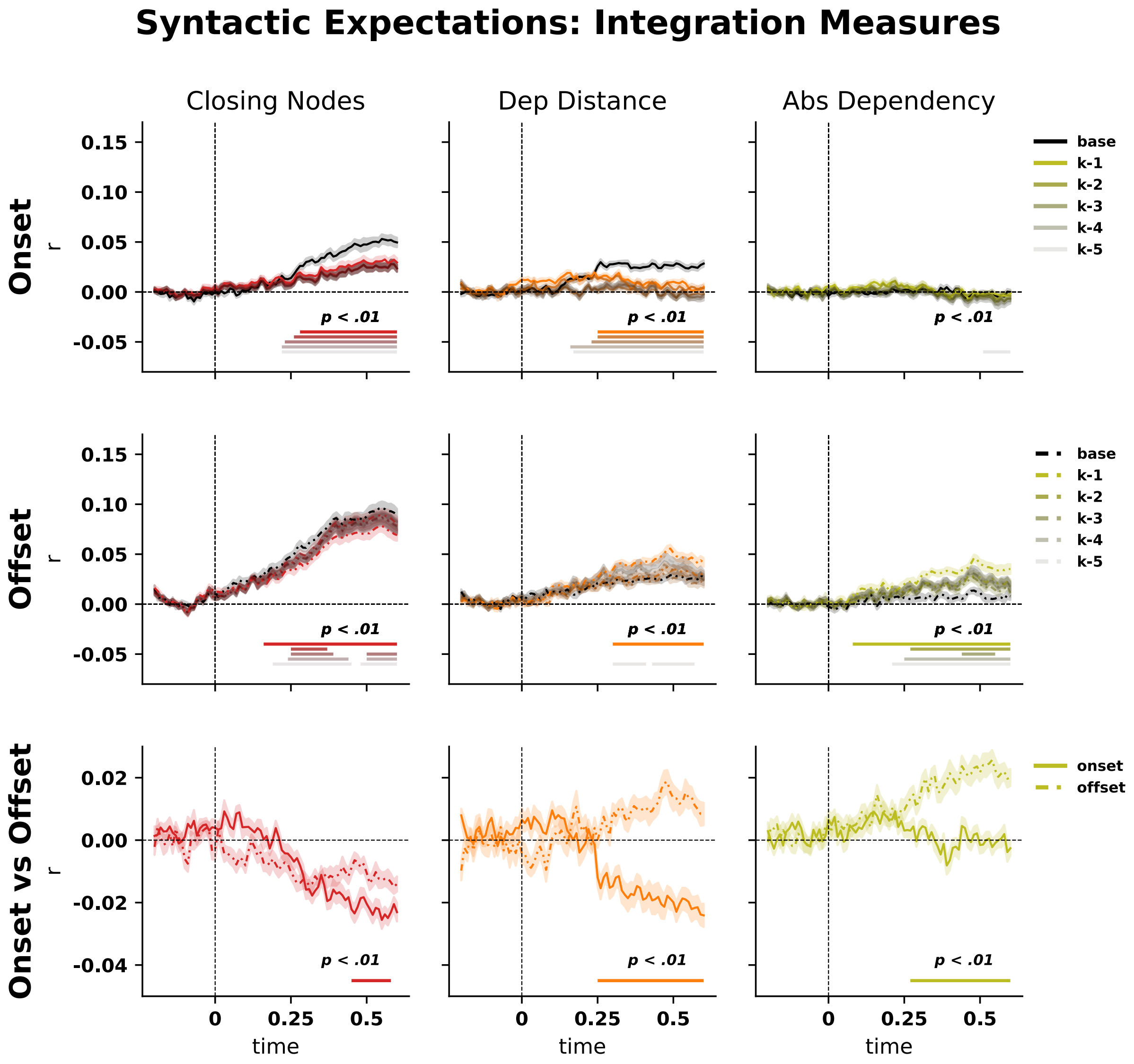


**Supplementary Fig. I 2** Partial correlation analysis of integration features. Before evaluating the decoding model's prediction we residualise our true variable and predicted variable and then compute the Pearson correlation. For integration measure we observe a reduction of the raw features, which allows context -3 to be also significantly better than baseline.
